# Lipidome profiling of human and equine neutrophil-derived extracellular vesicles and their potential contribution to the ensemble of synovial fluid-derived extracellular vesicles during joint inflammation

**DOI:** 10.1101/2023.11.17.567580

**Authors:** Laura Varela, Sanne Mol, Esther W. Taanman-Kueter, Sarah E. Ryan, Leonie S. Taams, Esther de Jong, P. René van Weeren, Chris H.A. van de Lest, Marca H.M. Wauben

**Author notes:** Corresponding authors: *E-mail address:* (M.H.M.Wauben).

## Abstract

The molecular signature of cell-derived extracellular vesicles (EVs) from synovial fluid (SF) offers valuable insights into the cells and molecular processes associated with joint disorders and can be exploited to define biomarkers. The signature of EVs is determined by cargo molecules and the lesser-studied lipid bilayer. We here investigated the lipidome of SF-EVs in inflamed joints derived from Rheumatoid Arthritis (RA) and Spondyloarthritis (SpA) patients, two autoimmune-driven joint diseases, and compared these signatures to the lipid profile of equine SF-EVs obtained during induced acute synovitis. Since neutrophils are primary SF-infiltrating cells during these inflammatory joint diseases, we also analyzed how inflammatory stimuli alter the lipidomic profile of human and equine neutrophil-derived EVs (nEVs) *in vitro* and how these signatures relate to the lipidome signatures of SF-EVs from inflamed joints. We identified neutrophil stimulation-intensity dependent changes in the lipidomic profile of nEVs with elevated presence of dihexosylceramide (lactosylceramide), phosphatidylserine, and phosphatidylethanolamine ether-linked lipid classes in human nEVs upon full neutrophil activation. In horses, levels of monohexosylceramide (glucosylceramide) increased instead of dihexosylceramide, indicating species-specific differences. The lipid profiles of RA and SpA SF-EVs were relatively similar and showed a relative resemblance with stimulated human nEVs. Similarly, the lipidome of equine synovitis-derived SF-EVs closer resembled the one of stimulated equine nEVs. Hence, lipidome profiling can provide insights into the contribution of nEVs to the heterogeneous pool of SF-EVs, deepening our understanding of inflammatory joint diseases and revealing molecular changes in joint homeostasis, which can lead to the development of more precise disease diagnosis and treatment strategies.

## 1. Introduction

Joint diseases represent a serious challenge in humans and animals, as they severely affect mobility and overall quality of life. In humans, these afflictions substantially strain the healthcare system Scott et al. [2022]; Leifer et al. [2022]. In animals, particularly horses, these disorders are a significant welfare concern that additionally causes significant economic losses to the equine industry Oke and McIlwraith [2010]. Virtually all joint diseases have an inflammatory component. The intensity of this inflammation, however, can vary widely: from the relatively mild and intermittent inflammation observed in osteoarthritis, which is a predominantly degenerative condition Sokolove and Lepus [2013], to the more aggressive and sometimes fulminant inflammation seen in septic arthritis de Grauw et al. [2009] or in autoimmune joint diseases such as rheumatoid arthritis (RA), and spondyloarthritis (SpA), which can cause severe and progressive tissue damage Merola et al. [2018]; Generali et al. [2018]. RA mainly affects peripheral joints, destroying cartilage and bone Lei et al. [2023], while SpA also affects the spine, skin, eyes, and gut Taitt and Balakrishnan [2023]. Joint disorders, in general, and subgroups of these, like rheumatic joint diseases, form a pathological spectrum rather than a collection of very distinct conditions. Detailed insights into the molecular endotype of neutrophil-associated arthropathies and the mechanistic backgrounds of these diseases may eventually aid in optimal diagnosis and targeted treatment.

Of the existing animal models for joint disease, the horse is acknowledged as being one of the best, given the strong resemblance with humans in regards to the histological appearance of the osteochondral unit, with its respective biochemical composition, the structure of the extracellular matrix of the articular cartilage, and the biomechanical characteristics of the tissue Malda et al. [2012]; McCoy [2015]; Moran et al. [2016]. Because of the high prevalence of joint diseases in horses and the heavy impact on the species that parallels the situation in humans, horses are considered not just a model species for human joint disease but also a target species in their own right. This prompted us to employ a well-established transient arthritis model in horses, which involves the intra-articular administration of lipopolysaccharide (LPS) to induce acute synovitis, for a comparative lipidome profiling study of synovial fluid (SF) derived EVs in inflammatory joint diseases de Grauw et al. [2009]; Cokelaere et al. [2018]; Varela et al. [2023].

Inflamed joints of human patients with RA, SpA, or septic arthritis and horses with acute synovitis exhibit a significant presence of infiltrated neutrophils de Grauw et al. [2009]; Loh and Lam [2022]; Coletto et al. [2023]; Boff et al. [2018]. Neutrophils serve as primary responders and are known to attract other immune cells to the site of action Loh and Lam [2022]. Apart from their classic role in eliminating pathogens, neutrophils have been shown to actively regulate adaptive immune responses. They can directly modulate T-cell and dendritic cell (DC) responses by activating DC maturation, stimulating CD4+ and CD8+ T cells, and modulating T-cell polarization Bert et al. [2023]; Groot Kormelink et al. [2018]. Their mode of action involves the release of compounds of highly cytotoxic granules and extracellular vesicles Bert et al. [2023], and uncontrolled activation of these mechanisms can harm healthy cells and the surrounding tissue.

Extracellular vesicles (EVs) are cell-derived lipid bilayer-enclosed particles from approximately 30 nm to 1 μm in size that are released in the extracellular environment and that actively participate in cell-to-cell communication Couch et al. [2021]. In recent years, EVs have garnered significant attention with regard to their roles in various physiological and pathological processes, including joint disorders. Synovial fluid (SF) contains significantly higher concentrations of EVs in individuals with inflammatory joint disease Varela et al. [2023]; Miao et al. [2022]; Born and Khachemoune [2023]; Foers et al. [2021]. Moreover, SF-derived EVs (SF-EVs) of oligoarticular juvenile idiopathic arthritis patients have a significantly altered protein profile in comparison to blood plasma-derived EVs of these patients and of healthy subjects Raggi et al. [2023]. Additionally, in RA patients, citrullinated proteins have been found to be associated with SF-EVs, and EV-associated miRNAs have been shown to regulate target genes associated with inflammation, e.g., STAT3, a key player in pro-inflammatory processes Foers et al. [2021]; Skriner et al. [2006]. Although the significance of SF-EVs in inflammatory joint pathologies is evident, their precise role in the disease processes remains largely undefined. The vast majority of molecular EV profiling is based on bulk analysis of the heterogeneous EV pool present within SF. Historically, proteomics and small RNA analysis of SF-EVs have been employed with the objective of revealing their cellular origins and disease biomarkers. However, lipids also play a pivotal role in EV biogenesis, secretion, and function Holopainen et al. [2019]. Notably, there is evidence for discernible alterations in the composition of the EV lipid bilayer, with a changing lipidome that seems to vary in accordance with the clinical state of the patient Phuyal et al. [2015]. Hence, alterations in the lipidome of SF-EVs during joint inflammation might provide valuable insights into the role of SF-EVs in joint diseases and might potentially serve as biomarkers. Importantly, in the LPS-induced acute synovitis equine model, we recently demonstrated, besides significant shifts in the quantities of SF-derived EVs, alterations in their lipidomic profiles during the acute inflammatory and resolving phases Varela et al. [2023].

To better understand the role of different EV subsets within the heterogenous SF-EV pool, we here explored whether a neutrophil-derived EV-associated lipid endotype could be identified. Hereto, we analyzed the impact of inflammatory stimuli on the lipidomic profiles of EVs sourced from primary human and equine neutrophils (nEVs) *in vitro*. Moreover, we analyzed the lipidomic profile of SF-EVs from human patients affected by RA or SpA and equine SF-EVs from the acute synovitis model while comparing these profiles to the lipidomes of neutrophil-derived EVs.

## Materials and Methods

### Specific procedures for human material

#### Neutrophil Isolation from Blood and Cell Culture Conditions

Blood samples (n=5) were obtained from healthy volunteers who provided informed consent. The blood collection protocol was approved by the institutional board of the Amsterdam Medical Centre (METC 2015_074). The blood was collected in sodium heparin tubes (Greiner Bio-One, Alphen a/d Rijn, The Netherlands). Neutrophils were isolated using a density gradient and erythrocyte lysis method, as described previously Mol et al. [2021]. The isolated neutrophils were suspended in IMDM (Iscove’s Modified Dulbecco’s Medium, Gibco, Thermo Fischer Scientific Inc., Waltham, United States) supplemented with EV-depleted heat-inactivated fetal bovine serum (FBS; Hyclone; Thermo Fischer Scientific Inc., Waltham, United States) and gentamycin (86 *μ*g/mL; Duchefa Biochemie B.V., Haarlem, The Netherlands), seeded in 24-well plates at 10^6^ cells/mL and cultured for 2 hours at 37 °C with or without activation stimuli, such as lipopolysaccharide (10 ng/mL, LPS; catalog no. L3024, from Escherichia Coli. O111:B4, Sigma-Aldrich, St. Louis, United States) or tumor necrosis factor-alpha (1 ng/mL, TNF-α; catalog no. 130-094-022, Miltenyi Biotec, Bergisch Gladbach, Germany) in combination with LPS. Supernatants were collected and centrifuged twice at 200 ×g for 10 min and twice at 500 ×g for 10 min at 4 °C to remove cells and debris before isolating the EVs.

### Neutrophil Flow Cytometric Analysis

The purity of neutrophils was assessed using flow cytometry, which was consistently above 97%. Briefly, the cells were washed twice in PBA (PBS-0.5% w/v BSA-0.05% w/v sodium azide) at 4 °C, then labeled with antibodies in PBA. The following antibodies were used: *α*CD15FITC (1:100; HI98), *α*CD16-PECy7 (1:1000; 3G8), *α*CD63-APC (1:100; H5C6), and *α*CD66b-PE (1:100; G10F5), all purchased from Biolegend (San Diego, United States). A FACSCanto instrument (BD Biosciences, San Jose, United States) was used to acquire 10,000 cells within the live gate, and FlowJo software version 10.7.1 (BD Biosciences) was employed for further analysis.

#### Collection of Synovial Fluid

During episodes of active arthritis, synovial fluid (SF) samples were collected through arthrocentesis from the knee joints of eight patients who provided informed consent. The synovial fluid collection was approved by the institutional board of the Amsterdam Medical Centre (METC 2018_04) and King’s College London through the PRISIM project by the Harrow Research Ethics Committee (17/LO/1940), IRAS ID 66106. Detailed information about the patients can be found in Supplementary Table S1. The samples were subjected to centrifugation at 650 ×g for 20 min at room temperature to remove cells, followed by centrifugation at 3000 ×g for 30 min at room temperature to remove any remaining cells and debris. The resulting cell-free SF was carefully collected and stored at -80 °C until needed for analysis.

### Specific procedures for equine material

#### Ethical Considerations

The *in vivo* experiments in this study were conducted under the Dutch Experiments on Animals Act and complied with the European Directive 2010/63/EU regarding the use of animals in research. The experiments were authorized by the Utrecht University Animal Ethics Committee (DEC), the Animal Welfare Body (IvD), and the Central Authority for Scientific Procedures on Animals (CCD) under application number AVD10800202013737.

#### Neutrophil Isolation from Blood and Cell Culture Conditions

Blood samples were collected from healthy adult warmblood horses and Shetland ponies in sodium heparin tubes (Greiner Bio-One, Alphen a/d Rijn, The Netherlands). Neutrophils were isolated using a density gradient (Lymphoprep®, StemCell Technologies, Vancouver, Canada) and underwent two rounds of hypotonic lysis to remove erythrocytes. The isolated neutrophils were then resuspended in IMDM supplemented with 10% EV-depleted heat-inactivated FBS (Gibco, Thermo Fischer Scientific Inc., Waltham, United States) and 100 *μ*g/mL Penicillin-Streptomycin (Gibco, Thermo Fischer Scientific Inc., Waltham, United States), seeded in 24-well plates at 106 cells/mL (24-well Costar® plates (Corning Inc., Corning, United States) and cultured for 2 hours at 37°C with or without activation stimuli, such as LPS (10 ng/mL, from Escherichia coli O55:B5; catalog no. L5418-2ML, Lot 118M4129V; St. Louis, United States) or equine TNF-*α* (1 ng/mL; catalog no. 1814-ET-025/CF, R&D Systems, Minneapolis, United States) in combination with LPS. Supernatants were collected and centrifuged twice at 200 ×g for 10 min and twice at 500 ×g for 10 min at 4 °C to remove cells and debris before isolating the EVs.

### Species-independent procedures

#### Extracellular Vesicle isolation

The isolation of extracellular vesicles (EVs) was performed following previously described methods Varela et al. [2023]; Boere et al. [2016]. Briefly, cell-free SF samples were incubated at 37 °C for 15 min with 20 *μ*L of hyaluronidase (HYase) (5 mg/mL; catalog no. H2126, Sigma-Aldrich, St. Louis, United States) to degrade hyaluronic acid. To eliminate protein aggregates, the SF was subsequently centrifuged at 1,000 ×g for 10 min at room temperature using an Eppendorf 5415R centrifuge with rotor F45-24-11. Next, supernatants from the SF samples and cellfree human and equine neutrophil cell culture supernatants were transferred to SW40 tubes (Beckman Coulter Inc., Brea, United States). They were subjected to centrifugation at 10,000 ×g for 35 min at 4 °C, followed by centrifugation at 100,000 ×g for 65 min using the SW40 Ti rotor in an Optima™ XPN-80 ultracentrifuge (Beckman Coulter Inc., Brea, United States). The EV pellets obtained from the 100,000 ×g spin of human neutrophil-derived EVs (Hu-nEVs) and SF-derived EVs (SF-EVs) were resuspended in 300 *μ*L of PBS with 0.1% BSA EV-depleted (previously depleted of EV and protein aggregates). The EV pellets from both 10,000 ×g and 100,000 ×g spins of equine neutrophil-derived EVs (Eq-nEVs) were mixed and resuspended in 100 *μ*L of PBS with 0.1% BSA EV-depleted.

For Hu-nEVs and SF-EVs, the EV pellets were mixed with 1.2 mL of a 2.5 M sucrose solution in PBS at the bottom of SW40 ultracentrifuge tubes. Subsequently, a series of fourteen sucrose solutions with decreasing molarity (ranging from 1.88 M to 0.4 M) were layered to create discontinuous sucrose gradients. The gradients were centrifuged at 40,000 rpm for 16 hours at 4°C using an Optima™ XPN-80 centrifuge with SW40 Ti rotor (average relative centrifugation force (RCF) of 202,048 ×g, maximum RCF of 284,570 ×g, κ-factor of 137; Beckman-Coulter, Fullerton, United States). After centrifugation, twelve fractions of 1 mL each were collected from the top (fraction 12) to the bottom (fraction 1), and the density of each fraction was determined using refractometry.

For Eq-nEVs, the EV pellets were mixed with 300 *μ*L of 60% Optiprep (StemCell Technologies, Vancouver, Canada) at the bottom of SW60 ultracentrifuge tubes. Subsequently, a series of eight Optiprep solutions with decreasing concentrations (ranging from 40% to 5%) were layered to create the gradients. The gradients were centrifuged at 45,000 rpm for 16 hours at 4 °C using an Optima™ XPN-80 centrifuge with SW60 Ti rotor (average RCF of 207,976 ×g, maximum RCF of 272,840 ×g, κ-factor of 81; Beckman-Coulter, Fullerton, United States). After centrifugation, eleven fractions of 333.3 *μ*L each were collected from the top (fraction 11) to the bottom (fraction 1), and the density of each fraction was determined using refractometry.

EV-enriched fractions (1.10-1.18 g/mL for sucrose gradients validated in Varela et al. [2023]; Mol et al. [2021] and 1.05-1.10 g/mL for Optiprep gradients validated in Boere et al. [2016]) were pooled. For Hu-nEVs and SF-EVs, the pooled fractions were loaded into SW32 ultracentrifuge tubes and pelleted by centrifugation for 95 minutes at 120,000 ×g at 4 °C (32,000 RPM; RCF average 127,755 ×g; RCF max 174,899 ×g; k-Factor 204). For the Eq-nEVs, the EV-enriched fractions were transferred to SW40 ultracentrifuge tubes and centrifuged for 95 minutes at 120,000 ×g at 4 °C (31,000 RPM; RCF average 121,355 ×g; RCF max 170,920 ×g; k-Factor 228). Subsequently, EV pellets were resuspended in 100 *μ*L PBS for immediate lipid extraction.

### Lipidomic analysis

#### Lipid Extraction

The extraction of lipids was carried out using a modified version of the Bligh& Dyer method Bligh and Dyer [1959]. Initially, 0.7 mL of fresh deionized water was combined with 100 *μ*L of the samples, 2 mL of methanol, and 1 mL of chloroform. The mixture was incubated for 20 min, followed by the addition of 2 mL of chloroform and 2 mL of deionized water. The resulting mixture was vortexed, allowing for the separation of the hydrophilic and hydrophobic phases through centrifugation at room temperature for 5 min at 2,000 ×g. The hydrophobic bottom phase was carefully transferred to a new conical glass tube. To ensure complete lipid collection, the extraction of the remaining sample (hydrophilic phase) was repeated with an additional 2 mL of chloroform. Subsequently, the samples were dried under nitrogen gas injection and stored in a nitrogen atmosphere at -20 °C.

#### Mass spectrometry lipidomics

The dried lipid pellets were reconstituted in 30 *μ*L of a chloroform/methanol mixture (1:1) and subjected to analysis following previously described methods Varela et al. [2023]; Jeucken et al. [2019].To ensure lipid quantification and serve as quality control, a sample composed of all individual samples in the same ratio, along with the SPLASH® Lipidomix® Mass Spec Standard (Avanti Polar Lipids, Inc., Alabaster, United States), was prepared and included in the mass spectrometry (MS) run. A hydrophilic interaction liquid chromatography (HILIC) column (2.6 *μ*m Kinetex HILIC 100, 50 x 4.6 mm; Phenomenex, Torrance, United States) was utilized, and 10 *μ*L of the lipid extract was loaded onto the column. The separation of lipid classes was achieved using a gradient elution on an Agilent 1290 InfinityII UPLC system (Agilent Technologies, Inc., Santa Clara, United States). Solvent A consisted of acetonitrile/acetone (9:1) with 0.1% formic acid, while solvent B was composed of acetonitrile/H2O (7:3) with 0.1% (v/v) formic acid and 10 mM ammonium formate (Gradient profile: from 0 to 1 minute, 100% A; from 1 to 3 min, 50% A + 50% B; from 3 to 5 min, 100% B, with a flow rate of 1 mL/min). The column was not re-equilibrated between injections, and the samples were directly injected.

The analysis was performed using a Fusion Orbitrap MS system (ThermoFisher Scientific, Waltham, United States) equipped with a heated electrospray ionization (HESI) source. The MS parameters included a negative ion spray voltage of 3.6 kV, an aux sheath gas flow rate of 54 Arb, a gas flow rate of 7 Arb, a sweep gas flow rate of 1 Arb, an ion transfer tube temperature of 350 °C, a vaporizer temperature of 450°C, and a scan range of 350-1100 m/z with a resolution of 120,000.

#### Lipid annotation

For the conversion of the RAW format to mzML, the open-source msconvert ProteoWizard software (ProteoWizard, Palo Alto, United States) was utilized Jeucken et al. [2019], with the selection of the “peakPicking filter vendor msLevel = 1-” parameter. LC/MS peak-picking, sample-grouping, and retention time-correction on the mzML files were performed using the XCMS package version 3.14.1 Smith et al. [2006]; Tautenhahn et al. [2008]; Benton et al. [2010], implemented in R free software version 4.1.2 Team [2020] (The R Foundation for Statistical Computing, Vienna, Austria). The LC/MS peaks (features) were annotated based on retention time, exact m/z-ratio, and the presence in at least 3 of all samples. An in silico lipid database was utilized for the annotation of MS peaks. Because the retention time is too close to the solvent front, ceramides (Cer) were ignored in this study. The selected features were also screened for isotope distribution and adducts. The original RAW files and the converted mzML mass spectrometry data have been deposited in the YODA repository of Utrecht University Varela and van de Lest [2023].

#### Data analysis

The lipidomics data were normalized based on the sum of total lipids per sample. This involved dividing each lipid value in a sample by the total sum of lipids in the same sample and multiplying it by 0.01, ensuring that the relative abundances summed up to 100. A minimum of three biological replicates were used for statistical analyses.

The data analysis was performed using the R software. For the principal component analysis (PCA), Pareto scaling was applied by dividing each variable by the square root of its standard deviation. Heatmap and cluster analysis were conducted on Spearman correlations among the 75 most abundant lipid species in all sample groups, with a set.seed value of two. These analyses were carried out using the R-package ComplexHeatmap 2.8.0 Gu et al. [2016]; Gu [2022].

The rank-based non-parametric Kruskal-Wallis test was employed to assess the differences between Hu-nEVs and SF-EVs, followed by multiple pairwise comparisons using Dunn’s test. Statistical significance was defined as a p-value less than 0.05. All Kruskal-Wallis and Dunn’s tests were conducted using GraphPad Prism 9 (GraphPad Software, Boston, United States).

## 2. Results

### Lipidomic Analysis of Human Neutrophil EVs Reveals Stimulation-Dependent Changes

To investigate how inflammatory stimuli alter the lipidome of primary human neutrophil-derived EVs (Hu-nEVs), we compared these EV-lipid profiles after activating neutrophils with a single stimulus (LPS), leading to partial neutrophil activation or with a dual stimulus (TNF*α* + LPS), inducing full neutrophil activation Mol et al. [2021]. Principal component analysis (PCA) shows that the differently stimulated neutrophils were separated in the plot having a sum variance of 71%, with EVs derived from dual stimulated neutrophils (TNF*α* + LPS) on the far left, followed by single stimulation (LPS) and unstimulated Hu-nEVs samples on the right (Figure 1A). These results indicate that the extent of neutrophil stimulation influenced the lipidome of Hu-nEVs.

**Figure 1:**
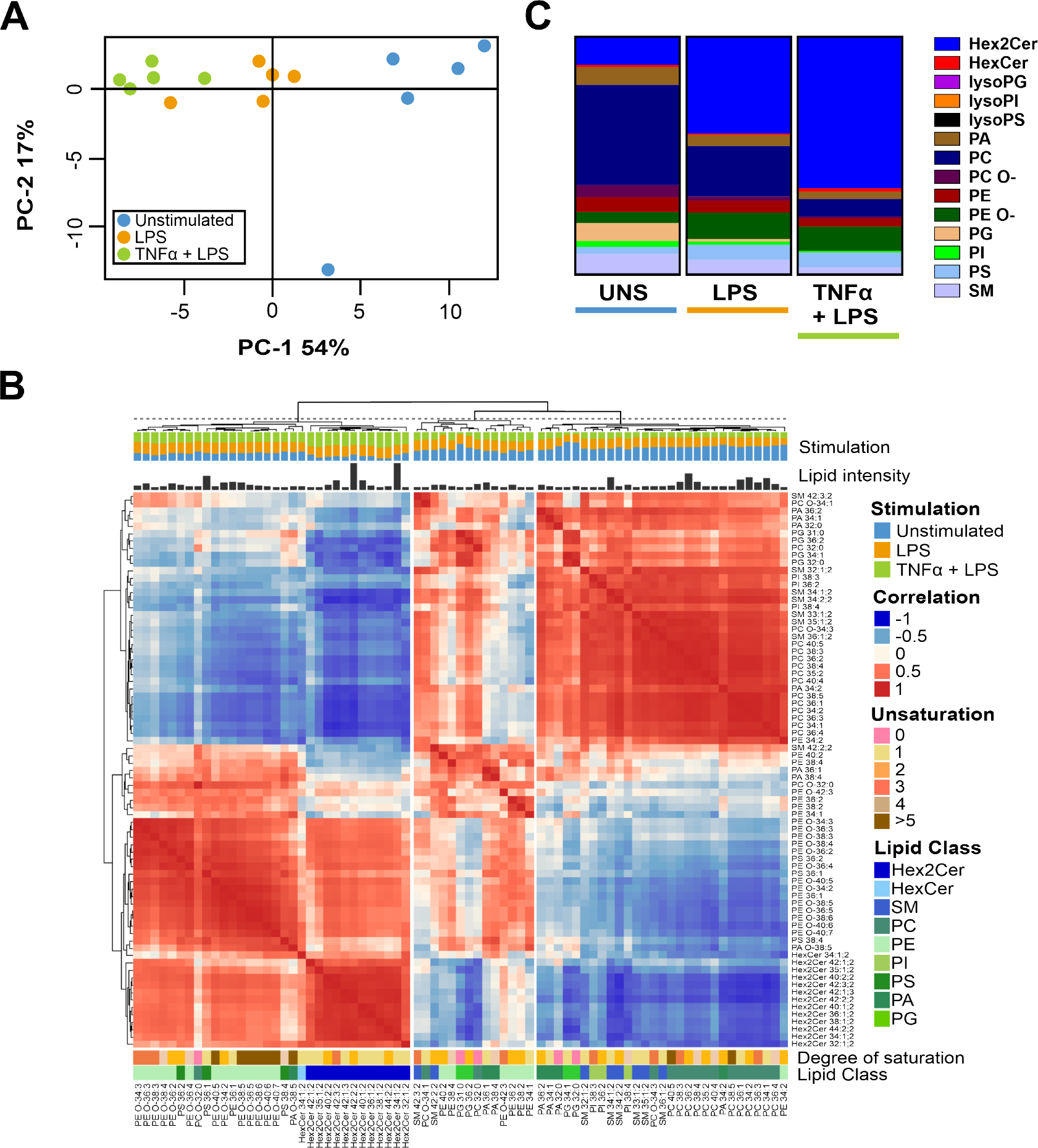
Lipidomic profiles of human neutrophil-derived EVs isolated from unstimulated primary neutrophils (Unstimulated), neutrophils partially stimulated with a single stimulus. **(i.e.**, **lipopolysaccharide (LPS)), and fully activated neutrophils stimulated with a dual stimulus (i.e**., **tumor necrosis factor-alpha and lipopolysaccharide (TNF***α***+LPS))**. Each group consisted of n=5 different blood donors. The EVs were isolated using a combination of differential ultracentrifugation steps at 10,000 ×g (10K) and 100,000 ×g (100K), followed by purification via sucrose density gradients. **A)** Principal Component Analysis (PCA) showing clear segregation of the different conditions. **B)** Combined heatmap (cluster dendrogram) of human neutrophil-derived EVs showing lipid-lipid Spearman correlations among the 75 most abundant lipid species across all EV sample groups. Partitioning Around Medoids, a centroidbased clustering approach sometimes referred to as K-Medoids, was used to determine the order of the lipids. Below the cluster dendrogram at the top of the figure, the group distribution is indicated (Stimulation), followed by the relative lipid intensity of each species and the heatmap. Beneath the heatmap, additional information is provided, including the degree of saturation, the lipid class of each lipid, and the respective annotation of each lipid species. **C)** Column plot illustrating the relative molar abundances for individual lipid classes. Abbreviated lipid classes: Hex2Cer (dihexosylceramide), HexCer (monohexosylceramide), Lyso (lyso-lipids), PG (phosphatidylglycerol), PS (phosphatidylserine), PI (phosphatidylinositol), PA (phosphatidic acid), O-(ether-linked lipids), PC (phosphatidylcholine), PE (phosphatidylethanolamine), and SM (sphingomyelin).

The heatmap (Figure 1B), based on a Spearman correlation between the 75 most abundant lipid species in all EV sample groups, revealed in the left branch notable correlations, such as the positive correlation between dihexosylceramide (Hex2Cer, most likely lactosylceramide: LacCer) and phosphatidylethanolamine (PE) ether-linked (O-) lipids, as well as between various phosphatidylserine (PS) species. These lipids were most abundant in Hu-nEVs derived from the fully activated neutrophils. Additionally, phosphatidylcholine (PC) lipids displayed stronger correlations with other PC lipids, leading to the formation of two distinct clusters in the right branch. These lipids were most prominent in the lipid profile of Hu-nEVs derived from unstimulated neutrophils. The lipid profile in relation to the lipid distributions (shown in the ‘stimulation’ annotation at the top of the heatmap) shows an evident progressive change with increasing neutrophil activation.

The relative molar abundances of individual lipid classes in Hu-nEVs are given in column plots (Figure 1C). The plot shows a stratified change in Hu-nEV lipid classes, primarily in Hex2Cer, PS, and PE ether-linked species. These key lipid classes demonstrated an increasing trend with increasing stimulation and neutrophil activation. In contrast, the quantities of PC, phosphatidylglycerol (PG), phosphatidic acid (PA), and sphingomyelin (SM) decreased progressively in EVs derived from fully activated neutrophils. Moreover, the comprehensive overview provided in Supplementary Figure 1) shows how neutrophil activation affected different lipid classes, such as the abrupt decrease of PG upon stimulation, which is barely present in EVs from fully activated neutrophils. Furthermore, the relative composition of individual lipid species within the major lipid classes did not change considerably upon neutrophil stimulation (Supplementary Figure1).

In summary, there was a discernible pattern in lipidome changes of Hu-nEVs derived from unstimulated to activated neutrophils, indicating a relationship between the activation status of neutrophils and the alteration of lipid classes of the released EVs.

### Comparative Lipidomic Characterization of synovial fluid derived-EVs in Neutrophil-Associated Arthropathies

To investigate whether lipidomic signatures of neutrophil-derived EVs can be identified in SF-derived EVs from patients with neutrophil-associated arthropathies, the lipidome of SF-EVs from RA and SpA patients was analyzed. The PCA of the SF samples from patients with RA and SpA showed a variance of 78% without a clear separation between the two diseases (Figure 2A). The correlation heatmap revealed three sections (Figure 2B). The first slice consisted of positive correlations among PS, some PE, some PE O-species, as well as a few PA and PI lipids. The second slice showed positive correlations between PC lipids, SM, and HexCer species, while the third section revealed positive associations between the remaining PE O-and Hex2Cer lipids. However, the overall lipidome did not display significant differences between the two disease groups. Also, a more detailed analysis of the lipid classes confirmed that the lipidomes of the samples from patients with RA and SpA (which are clinically classified as distinct disorders) were remarkably similar (Figure 2C, Supplementary Figure2). Similar to the Hu-nEVs profiles, the major lipid classes did not undergo significant changes in terms of lipid species composition, regardless of the type of arthropathy (Supplementary Figure2). These findings suggest that there are no substantial differences in the lipidomes of EVs in the synovial fluid among these two different arthropathies.

**Figure 2:**
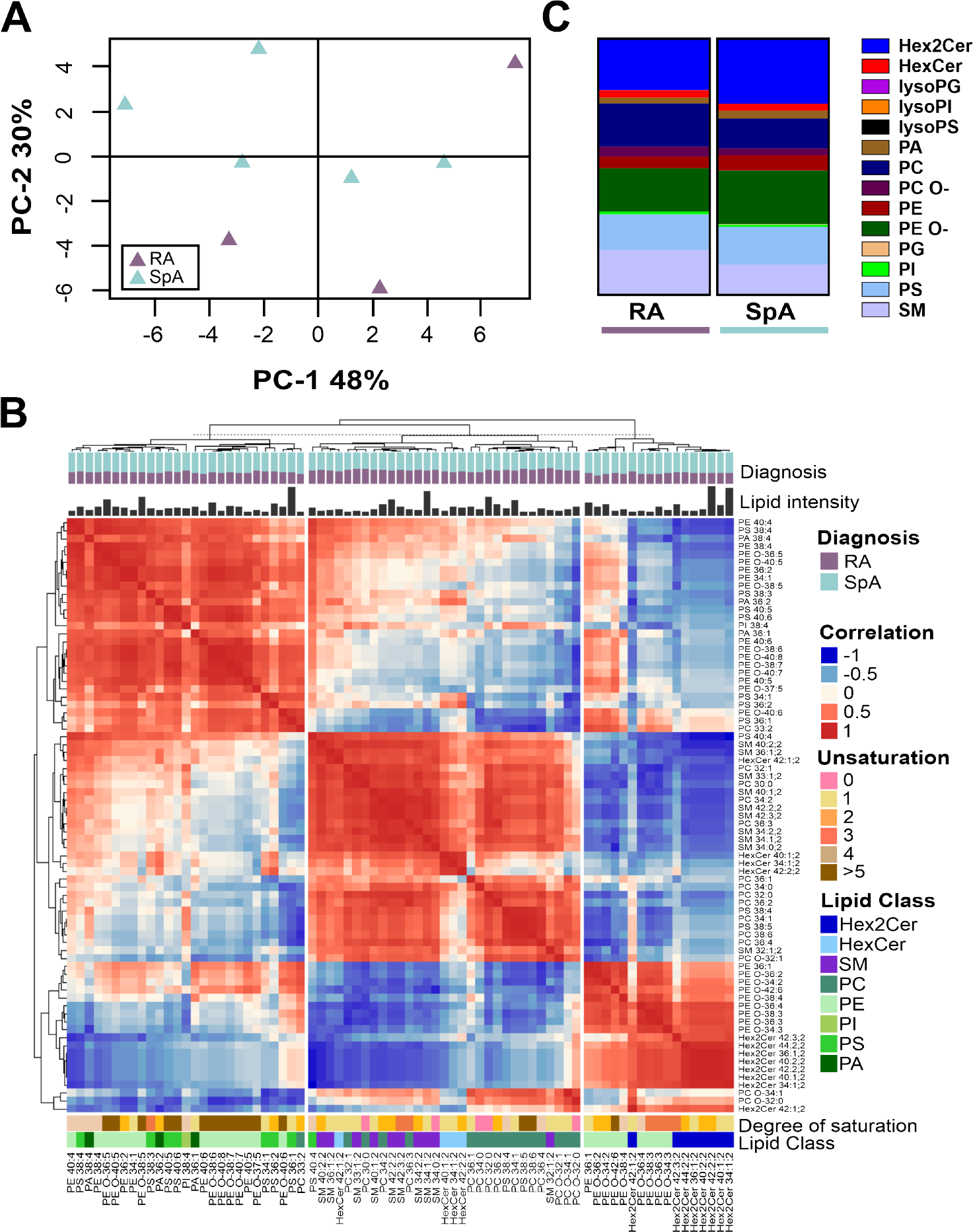
Lipidomic profile of human synovial fluid-derived EVs from rheumatoid arthritis (RA) and spondyloarthritis (SpA) patients. The RA group consisted of three patients (n=3), and the SpA group consisted of five patients (n=5). **A)** PCA conducted on the lipid profiles of the two arthropathies displayed no clustering of any of the groups. **B)** Combined heatmap (cluster dendrogram) of synovial fluid-derived EVs from RA and SpA patients showing lipid-lipid Spearman correlations among the 75 most abundant lipid species across all EV sample groups. Partitioning Around Medoids, a centroid-based clustering approach sometimes referred to as K-Medoids, was used to determine the order of the lipids. Below the cluster dendrogram at the top of the figure, the group distribution is indicated (Stimulation), followed by the relative lipid intensity of each species and the heatmap. Beneath the heatmap, additional information is provided, including the degree of saturation, the lipid class of each lipid, and the respective annotation of each lipid species. **C)** Lipid classes of SF-EVs of RA and SpA SF-EVs, showing the relative size of each lipid class (%) on the y-axis. Abbreviations: Hex2Cer (dihexosylceramide), HexCer (monohexosylceramide), Lyso ( lyso-lipids), PG (phosphatidylglycerol), PS (phosphatidylserine), PI (phosphatidylinositol), PA (phosphatidic acid), O-(ether-linked lipids), PC (phosphatidylcholine), PE (phosphatidylethanolamine), and SM (sphingomyelin).

### Comparison of the lipidomes of human neutrophil-derived EVs with synovial fluid-derived EVs in Neutrophil-Associated Arthropathies

Based on the presence of specific protein markers, it has been indicated that neutrophil-derived EVs are highly abundant, at least in SF from RA patients Foers et al. [2020]. In this study, we performed a comparative lipidomic analysis of SF-EVs from RA and SpA patients and the lipidome of hu-nEVs, with the objective of ascertaining whether lipid profiles specific to neutrophil-EVs can be identified in the SF of these patients. The PCA analysis revealed that the SF-EVs from the two arthropathies clustered together at the bottom of the PCA quadrant (Figure 3A). Notably, the closest clusters to these samples were that of the Hu-nEVs derived from single-stimulated neutrophils, followed by the Hu-nEVs from fully activated neutrophils. The key lipids that caused the greatest variance can be divided into three groups, including species from the Hex2Cer, PC, SM, PS, and ether-linked PE lipid classes (Figure 3B). In Supplementary Figure 3, the individual lipid species that defined the loading plot of the PCA shown in Figure 3B are indicated. The Hex2Cer 42:2;2 and Hex2Cer 34:1;2 lipid species were significantly increased in Hu-nEVs from fully activated and single-stimulated neutrophils compared to unstimulated Hu-nEVs and SF-EVs (from respectively RA and SpA or SpA patients). While in SF-EVs, Hex2Cer 42:2;2 and Hex2Cer 34:1;2 were significantly increased compared to EVs derived from unstimulated neutrophils (Supplementary Figure 3, Figure 3C). Furthermore, PS 36:1 exhibited a significant upsurge in Hu-nEVs derived from fully activated and single-stimulated neutrophils, as well as SF-EVs from RA and SpA, relative to unstimulated neutrophils. PC, conversely, is diminished in Hu-nEVs from stimulated neutrophils and SF-EVs from patients with neutrophil-associated arthropathies compared to unstimulated neutrophil-derived EVs. As such, a significant decrease was shown in lipid species PC 36:2, PC 34:2, PC 34:1, and PC 36:1 in, respectively, SF-EVs and Hu-nEVs derived from both fully activated and single-stimulated neutrophils, as compared to Hu-nEVs from unstimulated counterparts (Supplementary Figure 3, Figure 3C). In contrast to the decrease in the previously mentioned PC species in Hu-nEVs from stimulated neutrophils and SF-EVs, the relative contribution of lipid species PC 32:0 was significantly increased in SF-EVs derived from both RA and SpA patients as compared to all Hu-nEV conditions (Supplementary Figure 3, Figure 3C). A similar relative increase in SF-EVs was observed for the PE-ether linked lipids PE-O 38:5 and PE-O 36:5, and SM lipids 42:2;2, and SM 34:1;2.

**Figure 3:**
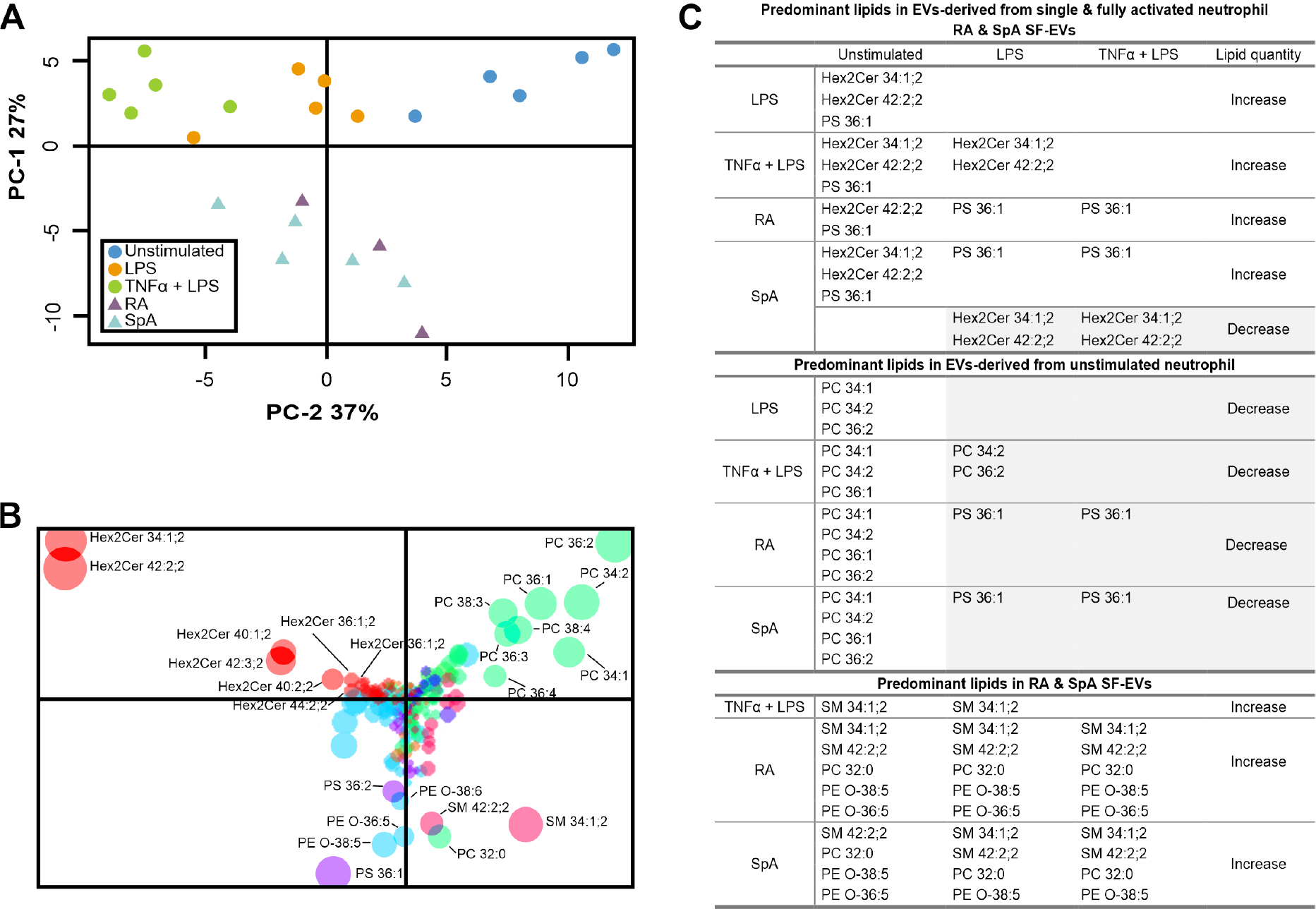
Comparative Analysis of Lipid Profiles in SF-EVs from RA, SpA, and EVs from differently stimulated human neutrophils. **A**) Overview of PCA sample distribution. The PCA score plot of principal component-1 (PC-1) and PC-2 demonstrates the location of the SF-EV samples in comparison to the Hu-nEVs. **B)** Corresponding PCA loading plot. The PCA loading plot indicates individual lipids identified in the samples. The size of the dots represents the log of the summed intensity of each lipid. **C)** Comparative analysis of significant lipids from panel B. For a comprehensive lipid description, see Supplementary Figure 3. A decrease in lipid intensity when comparing conditions (column vs. row, exemplified by LPS vs. Unstimulated and TNF*α* + LPS vs. Unstimulated) is highlighted in light gray, while increases are left uncolored. Abbreviations: Hex2Cer (dihexosylceramide; lactosylceramide), O-(ether-linked lipids), PC (ester-linked phosphatidylcholine), PE (phosphatidylethanolamine), PS (phosphatidylserine), SM (sphingomyelin), RA (rheumatoid arthritis), SpA (spondyloarthritis), TNF*α* (tumor necrosis factor-*α*) and lipopolysaccharide (LPS).

The latter findings suggest contributions from EVs of other cellular origins than Hu-nEVs in the total ensemble of SF-EVs from the patients.

### Comparison of the lipidomes of equine neutrophil-derived EVs with equine synovial fluid EVs from healthy and acute LPS-induced synovitis joints

Recently, we observed distinct changes in the lipid composition of equine SF-EVs from healthy joints compared to joints with lipopolysaccharide (LPS)-induced synovitis (i.e., during acute inflammation), where monohexosylceramides (HexCer, also known as GlcCer: glucosylceramide or GalCer: galactosylceramide) were significantly elevated and PS, PC, and SM were reduced in SF-EVs Varela et al. [2023]. Since this equine model is characterized by a strong neutrophil influx in the acute phase, we aimed to analyze whether the observed changes in the lipid signatures of SF-EVs could be attributed to EVs derived from equine neutrophils. Hereto, we investigated EVs released by primary equine neutrophils (Eq-nEVs) that were either unstimulated or dual stimulated with TNF*α* + LPS. The Principal component analysis (PCA) shows an explained variance of 85%, which resulted in a partial separation between Eq-nEVs derived from unstimulated or fully activated neutrophils (Figure 4A).

**Figure 4:**
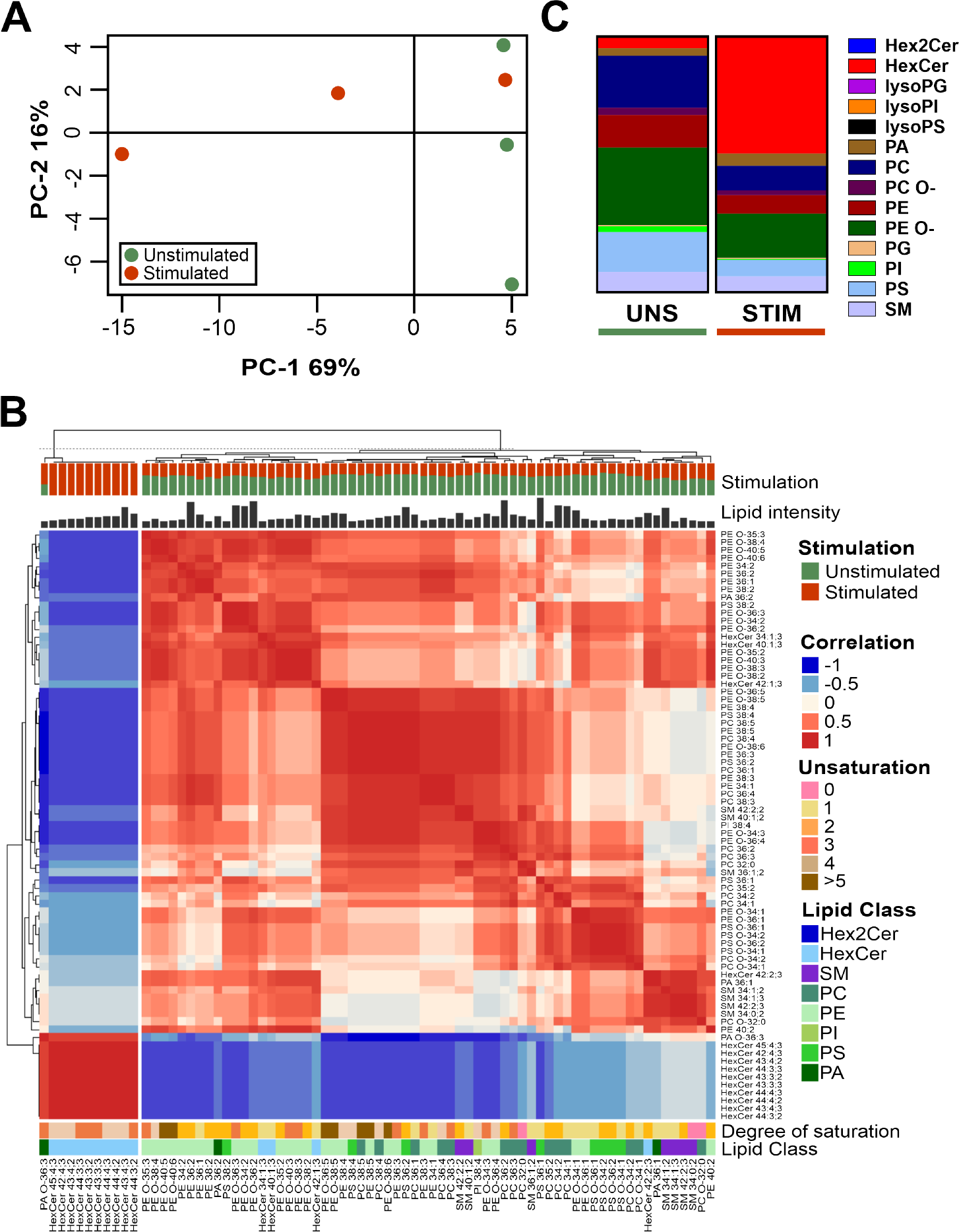
Lipidomic profiles of equine neutrophil-derived EVs isolated from unstimulated primary neutrophils (Unstimulated) and fully activated neutrophils stimulated with a dual stimulus. (**i.e.**, **tumor necrosis factor-alpha and lipopolysaccharide (TNF***α***+LPS))**. Each group consisted of n=3 different blood donors. **A)** Principal Component Analysis (PCA) showing clear segregation of the different conditions. **B)** Combined heatmap (cluster dendrogram) of equine neutrophil-derived EVs showing lipid-lipid Spearman correlations among the 75 most abundant lipid species across all EV sample groups. Partitioning Around Medoids, a centroid-based clustering approach sometimes referred to as K-Medoids, was used to determine the order of the lipids. Below the cluster dendrogram at the top of the figure, the group distribution is indicated (Stimulation), followed by the relative lipid intensity of each species and the heatmap. Beneath the heatmap, additional information is provided, including the degree of saturation, the lipid class of each lipid, and the respective annotation of each lipid species. **C)** Column plot showing the relative molar abundances for individual lipid classes of Eq-nEVs. Abbreviated lipid classes: Hex2Cer (dihexosylceramide), HexCer (monohexosylceramide), Lyso (lyso-lipids), PG (phosphatidylglycerol), PS (phosphatidylserine), PI (phosphatidylinositol), PA (phosphatidic acid), O-(ether-linked lipids), PC (phosphatidylcholine), PE (phosphatidylethanolamine), and SM (sphingomyelin).

The correlation heatmap showed a strong cluster of HexCer in the left branch of the dendrogram that was exclusively associated with Eq-nEVs derived from fully activated neutrophils (Figure 4B). The other lipid classes clustered together and were more prominent in Eq-nEVs samples derived from unstimulated neutrophils. When evaluating the specific lipid classes (Figure 4C, Supplementary Figure 4), it became apparent that in contrast to Hu-nEVs, the relative amount of Hex2Cer in the entire lipidome is low. Surprisingly, a strong increase in HexCer levels in Eq-nEV samples derived from fully activated neutrophils was observed. This change resulted in a relative reduction of all other lipid classes except for PA (Figure 4, Supplementary Figure 4). When normalizing the lipid species within each class, it became evident that most classes did not undergo significant changes in their lipid profiles, except for Hex2Cer and HexCer. Remarkably, in the Hex2Cer class, the contribution of Hex2Cer 42:2;2 and Hex2Cer 34:1;2, the two dominant Hex2Cer types in Hu-nEVs decreased in Eq-nEVs derived from activated neutrophils in which Hex2Cer 33:2;2 and Hex2Cer 32:2;2, both low abundant in Hu-nEVs, dominate the Hex2Cer class. The profile of HexCer showed that all the species present in unstimulated neutrophil-derived Eq-nEVs decreased relatively upon stimulation, as several new long-chain lipid species appeared (from HexCer 42:4;3 to HexCer45:5;3).

Subsequently, a comparative analysis of the lipid profiles of Eq-nEVs derived from unstimulated and fully activated primary equine neutrophils and the previously described profiles of SF-EVs from healthy equine joints and those afflicted by acute LPS-induced synovitis was performed Varela et al. [2023]. Comparison of the relative contribution of individual lipid species to a lipid class showed that elevated HexCer lipid species, i.e., HexCer 34:1;3, Hex-Cer 38:1;3, HexCer 40:1;3 and HexCer 42:2;3, that were paramount in the transition from healthy to inflamed state in the LPS-synovitis model were also identified in Eq-nEVs (Table 1). Remarkably, despite a strong relative increase in the HexCer lipid class in Eq-nEVs upon full neutrophil activation (Figure 4, Supplementary Figure 4), these Eq-nEVs showed a discernible decrease in the relative contribution of HexCer 34:1;3, HexCer 38:1;3, HexCer 40:1;3 and HexCer 42:2;3 caused by the diversification of the HexCer lipid class of Eq-nEVs derived from fully activated equine neutrophils (Table 1, Supplementary Figure 4). For the SM lipid class, SF-EVs from the LPS model presented augmented levels of SM 42:2;3, SM 42:1;3, SM 34:0;3, and SM 34:1;3 during acute synovitis, a trend that was mirrored within the Eq-nEVs from activated neutrophils (Table 1) Concerning the PE lipid class, SF-EVs sourced from acute synovitis (and not from healthy SF-EVs) illustrated an upsurge in the PE class, predominated by ether-linked PE species Varela et al. [2023]. Noteworthy species included PE 38:2, PE O-34:2, PE O-36:2, PE O-36:3, PE O-38:2, PE O-38:3, and PE O-40:3 (Table 1). These species were also present in Eq-nEVs, but the quantities remained reasonably consistent (relative to the respective complete lipidome) irrespective of whether the neutrophils were stimulated or unstimulated. Similarly to ether-linked PE, PC O-34:1, PC O-34:3, and PC O-38:3 had relatively constant levels in nEVs from unstimulated and fully activated neutrophils compared to SF-EVs, whereas during acute synovitis, a relative increase of these ether-linked PC species was observed (Table 1). Only PC O-32:0 showed a relative increase in SF-EVs during acute synovitis and equine nEVs from activated neutrophils.

**Table 1:** Comparison of the relative quantities (%) of lipid species (mean ± SD) that predominantly increase in acutely inflamed equine SF-EVs vs. healthy SF-EVs Varela et al. [2023] with equine nEVs derived from unstimulated and fully activated primary neutrophils.

| Lipid species | Healthy - SF-EVs | Acute synovitis - SF-EVs | UNS Eq-nEVs | STIM Eq-EVs |
| --- | --- | --- | --- | --- |
| HexCer 34:1;3 | 4.21 ± 0.72 | 11.51 ± 0.92 | 6.54 ± 5.34 | 3.01 ± 4.76 |
| HexCer 38:1;3 | 1.01 ± 0.92 | 5.58 ± 1.51 | 3.02 ± 2.42 | 1.8 ± 2.9 |
| HexCer 40:1;3 | 6.72 ± 1.07 | 16.94 ± 2.23 | 5.88 ± 3.69 | 3.62 ± 5.31 |
| HexCer 42:2;3 | 6.57 ± 3.03 | 47.56 ± 0.33 | 22.23 ± 8.43 | 8.21 ± 10.94 |
| PE 38:2 | 0.16 ± 0.06 | 2.51 ± 1.98 | 2.24 ± 1.0 | 2.85 ± 2.4 |
| PE O-34:2 | 2.93 ± 0.45 | 13.44 ± 0.75 | 10.56 ± 2.38 | 11.49 ± 1.58 |
| PE O-36:2 | 1.33 ± 0.04 | 16.79 ± 0.99 | 14.68 ± 6.97 | 17.82 ± 5.61 |
| PE O-36:3 | 1.25 ± 0.02 | 8.12 ± 0.41 | 13.49 ± 3.33 | 12.4 ± 3.37 |
| PE O-38:2 | 0.23 ± 0.01 | 6.33 ± 0.26 | 2.38 ± 1.06 | 4.77 ± 2.6 |
| PE O-38:3 | 0.22 ± 0.03 | 6.36 ± 0.27 | 3.06 ± 1.59 | 4.47 ± 2.17 |
| PE O-40:3 | 0.11 ± 0.01 | 3.44 ± 0.2 | 1.45 ± 0.85 | 2.06 ± 1.19 |
| SM 42:2;3 | 0.43 ± 0.03 | 22.32 ± 2.32 | 5.86 ± 3.77 | 9.31 ± 3.79 |
| SM 42:1;3 | 0.21 ± 0.06 | 6.55 ± 2.82 | 2.38 ± 1.43 | 3.25 ± 1.64 |
| SM 34:0;3 | 0.12 ± 0.08 | 7.03 ± 5.6 | 0.45 ± 10.07 | 0.97 ± 6.96 |
| SM 34:1;3 | 0.02 ± 0.01 | 5.37 ± 0.47 | 2.41 ± 0.37 | 4.18 ± 0.35 |
| PC O-32:0 | 0.81 ± 0 | 6.97 ± 0.79 | 7.06 ± 2.9 | 11.29 ± 6.63 |
| PC O-34:1 | 1.49 ± 0.19 | 7.54 ± 0.31 | 32.71 ± 2.88 | 31.27 ± 13.26 |
| PC O-34:3 | 0.23 ± 0.12 | 2.9 ± 0.35 | 5.34 ± 1.11 | 4.74 ± 2.74 |
| PC O-38:3 | 0.01 ± 0 | 2.74 ± 0.02 | 1 ± 1.04 | 0.98 ± 1.25 |
| PS 36:1 | 51.35 ± 3.27 | 79.97 ± 0.11 | 40.48 ± 1.71 | 48.73 ± 8.54 |
| PS 36:2 | 48.65 ± 3.27 | 20.03 ± 0.11 | 23.08 ± 11.03 | 21.58 ± 9.35 |

Although the PS class demonstrated notable lipid diversity in the Eq-nEVs (Supplementary Figure 4), the lipid species PS 36:1 and PS 36:2 dominated in both Eq-nEVs from unstimulated and stimulated neutrophils, as well as in SF-EVs of both healthy joints and during acute synovitis Varela et al. [2023]. During acute synovitis, the relative contribution of PS 36:1 in SF-EVs increases. This trend is also observed in Eq-nEVs from stimulated neutrophils, albeit the change in the ratio of the two PS species is less pronounced.

In conclusion, similar to what was observed with RA and SpA SF-EVs compared to the Hu-nEVs, our comprehensive analysis and comparison unveiled lipid signatures of Eq-nEV in SF-EVs of the LPS-induced synovitis model. However, besides these similarities, intricate lipidome variations between Eq-nEVs and equine SF-EVs were identified, indicative of other EV subsets present in the heterogeneous SF-EV pool.

## 3. Discussion

In this study, we investigated the impact of *in vitro* neutrophil activation on the lipid composition of human and horse neutrophil-derived-EVs (nEVs) in order to define lipid signatures of nEVs and to relate these signatures to the lipid profiles of SF-EVs in neutrophil-associated arthropathies. Correlation analysis demonstrated significant changes in different lipid species of neutrophil-derived EVs upon neutrophil activation. Hence, neutrophil activation influences the lipid signature of neutrophil-derived EVs.

Previously, we showed that *in vitro* stimulation of healthy blood donor-derived neutrophils resulted in enhanced release of EVs, which was most pronounced upon dual stimulation Mol et al. [2021]. We here show that neutrophil activation substantially influenced the lipidome of Hu-nEVs, with lipid classes Hex2Cer and PS and ether-linked PE species showing a positive correlation with the degree of neutrophil stimulation, while lipid classes PC, PG, PA, and SM showed a negative correlation.

The lipid profiling of SF-EVs from the neutrophil-related arthropathies RA and SpA in this study demonstrated rather similar lipid profiles, with most lipid classes remaining consistent across both arthropathies. Comparison of SF-EV profiles with the profile of Hu-nEVs derived from unstimulated human neutrophils showed upregulation of Hex2Cer lipid species (i.e., Hex2Cer 42:2;2 and Hex2Cer 34:1;2) and PS 36:1, and downregulation of certain PC species (i.e., PC36:2, PC34:2 and PC34:1) as similarly observed in the comparison of unstimulated versus single and dual stimulated Hu-nEVs. Interestingly, the relative concentrations of the lipid classes Hex2Cer, PA, PE, PI, and PC aligned more closely with Hu-nEVs derived from single-stimulated than from dual-stimulated neutrophils. Hence, these results hint towards a milder neutrophil activation state *in vivo* compared to the pronounced effects observed after dual stimulation of neutrophils *in vitro*. Recently, it has been suggested that hyaluronic acid possesses an inhibitory effect on neutrophil activation in synovial fluid of SpA and RA patients Mol et al. [2023]. It is conceivable that *in vivo* hyaluronic acid plays a protective role, potentially mitigating possible excessive neutrophil activation. As such, this raises the question of whether the dual stimulation as studied *in vitro* truly replicates *in vivo* conditions or merely represents an ‘over-activated’ state in comparison to neutrophils present in SF during neutrophil-associated arthropathies. Further investigations are essential to ascertain the significance of these findings.

Importantly, the normalized lipid intensities of the individual lipid species, SM 42:2;2, SM 34:1;2, PS 36:1, PC 32:0, PE O-36:5, and PE O-38:5 in SF-EVs, displayed a marked increase compared to Hu-nEVs. These findings suggest that EV subpopulations in SF from RA and SpA could, apart from neutrophils, derive from other cell types linked directly to both arthropathies Povoleri et al. [2023]; Roberts et al. [2015]; Skougaard et al. [2022]. The increased levels of SM and PE O-in SF-EVs correlate well with higher levels in T cells, monocytes, B cells, and NK cells as compared to neutrophils, while the ratio (approximately 75%) of the two primary PS lipid species (PS 32:1 and PS 32:2) in SF-EVs resembles more closely the ratio of those PS species as described for B cells, T cells, NK cells, and monocytes Pernes et al. [2023] Also, the higher abundance of HexCer in SF-EVs from RA and SpA patients, as compared to nEVs, might originate from human T cells or monocytes-derived EVs since HexCer is not predominantly present in human neutrophils Pernes et al. [2023]; Alarcon-Barrera et al. [2020]. In T cells, gangliosides are overexpressed Nakayama et al. [2018]; Zhang et al. [2019]. Unfortunately, gangliosides were not measured in this study due to their higher mass, which was outside the analyzed range of our optimized settings for the detection of glycerophospholipids. However, the expression of GM3 ganglioside (monosialodihexosylganglioside) has been demonstrated to be higher in the synovia from RA patients compared to osteoarthritis patients Nakayama et al. [2018]. The unveiled complexity of the ensemble of SF EVs derived from inflamed joints raises questions about the exact contributions of EVs from different cellular origins to the SF-EV pool and the potential roles of these EV subsets in RA and SpA pathogenesis and urges for further SF EV subset analysis.

Also, the lipidome analysis of Eq-nEVs revealed marked changes depending on the neutrophil activation status, with significantly elevated levels of HexCer species upon dual stimulation. While Eq-nEVs from *in vitro* fully activated horse blood-derived neutrophils have nearly 50% of their lipidome composed of HexCer, SF-EVs from LPS-induced synovitis at the pinnacle of the inflammatory reaction contain around 15% HexCer of their total lipidome Varela et al. [2023]. Although the exact proportion cannot be determined based on our data, a substantial amount of the nEVs observed in SF might originate from activated neutrophils. In this same line, PC levels show a similar change, i.e., a decrease in the relative lipidome contribution after neutrophil stimulation and in SF-EVs at 5 hours post-injection of LPS *in vivo* Varela et al. [2023]. However, in the equine model studies, we observed undeniable differences between the lipidomes of stimulated Eq-nEVs and SF-EVs during LPS-induced synovitis, with a remarkable resemblance to the findings in SF-EVs from RA and SpA patients. For instance, SF-EVs from induced synovitis exhibit substantially more elevated SM levels than those seen in Eq-nEVs. Similarly, PE (both ether link and ester-link) levels decrease in Eq-nEVs from fully activated neutrophils, while PE (combined, i.e., esther-link and ether-link) classes augment in SF-Evs of acute synovitis Varela et al. [2023]. As mentioned before, such disparities can be linked to EVs from different cellular origins, contributing to the ensemble of SF-EVs.

Interestingly, we discovered a species-specific difference between the lipidomic profiles of Hu-nEVs and Eq-nEVs derived from stimulated neutrophils in terms of upregulated Hex2Cer and HexCer levels, respectively. The chemical distinction between these two sphingolipid classes lies solely in an extra hexose (most likely a galactose). Previously, it has been hypothesized that HexCer plays a crucial role in the production of TNF*α* and prostaglandins, both wellknown inflammatory mediators Qu et al. [2018]. Similarly, Hex2Cer has been proven to drive inflammation through the production of prostaglandins and an increase in the expression of CD11B/CD18 Chatterjee et al. [2021]. Therefore, nEVs enriched in HexCer or Hex2Cer present in the joint space during inflammatory episodes of RA, SpA, or LPS-induced synovitis might favor the production of TNF*α* and inflammatory lipid mediators and, as such, might be involved in enhancing and even perpetuating inflammatory responses. On the contrary, Hex2Cer may have protective effects, as Hex2Cer can bind several pathogens Nakayama et al. [2018], and LacCer (the major Hex2Cer) has been demonstrated to be directly involved in phagocytosis and the generation of superoxide Iwabuchi and Nagaoka [2002]; Iwabuchi et al. [2015]. In line with this, antibacterial properties have been demonstrated for subpopulations of activated neutrophil-derived EVs Lőrincz et al. [2019]. Unfortunately, the roles of HexCer have not been investigated to the same extent.

The distinctive lipidomic profile of nEVs originating from fully activated neutrophils in both human and equine species, in comparison to nEVs derived from unstimulated neutrophils, displays a noteworthy compositional shift. This alteration is expected, given the dynamic nature of neutrophils during activation, such as massive degranulation and NETosis, a unique process of neutrophils. The increased relative contribution of sphingolipids to the lipidome and the simultaneous decrease in PC within the lipid profile is probably stabilized by cholesterol, which is known to be particularly enriched in EVs Skotland et al. [2023]. It is essential, however, to recognize that the profiling of nEVs covers a diverse pool of different EV subsets, including exosomes, which are derived from the endosomal route, and ectosomes originating from the plasma membrane. Consequently, variations in EV size, shape, and biogenesis routes may be inherent to these distinct subsets. Indeed, previous studies have highlighted the heterogeneity in sizes and shapes of nEVs Lőrincz et al. [2014]. In addition, the influence of environmental factors during biogenesis has been demonstrated to affect the overall composition of the EV population, thereby impacting aspects of morphology, protein abundance, and, subsequently, functional attributes Kolonics et al. [2021]; Lőrincz et al. [2015].

The lipidomic composition of nEVs from activated neutrophils has been indicative of the presence of ectosomes (e.g., microvesicles or apoptotic bodies) Hess et al. [1999]. Additionally, the diverse sizes and shapes observed in nEVs suggest potential heterogeneity in biogenesis routes. Importantly, the annexin V binding capacity of neutrophil-derived ectosomes implies phosphatidylserine (PS) exposure, a characteristic feature of conventional apoptotic bodies Lőrincz et al. [2015]. Furthermore, the presence of EV-associated Lselectin, a membrane protein diminished of the cell surface upon neutrophil stimulation, further underscores the presence of plasma membrane-derived EVs Gasser et al. [2003]. Overall, the nEV population, composed of both exosomes and ectosomes, may change substantially upon neutrophil activation, after which the resultant ectosome population may also originate from various cell death processes, including NETosis, apoptosis, or ferroptosis Vorobjeva and Chernyak [2020].

In the current study, we did not evaluate the lipidome of neutrophils at the cellular level. However, it has been demonstrated for human neutrophils that Hex2Cer is the primary sphingolipid, comprising up to 70% of the cell profile Chatterjee et al. [2021]; Symington [1989]. Importantly, differences in Hex2Cer enrichment between species have been described, albeit data on horse neutrophils are lacking Pernes et al. [2023]. In mouse neutrophils, no aberrant enrichment of Hex2Cer has been observed but rather a combination of SM, HexCer, and ceramide Pernes et al. [2023]. Nevertheless, a modest amount of Hex2Cer is discernible in the mouse neutrophil lipidome. Equally, we here show that equine nEVs also contain traces of Hex2Cer besides a strong enrichment of HexCer. The reason for the preference for HexCer or Hex2Cer enrichment in different species is not known and warrants deeper investigation.

In conclusion, our study offers valuable novel insights into the intricate relationship between neutrophil stimulation and nEV lipidome changes. Further research is required to validate these findings and to unravel the interplay of various cellular entities and their EVs in the inflammatory processes in joint diseases. Overall, our findings deepen our understanding of the lipid dynamics in neutrophil-associated arthropathies and contribute to our knowledge of the changes in SF-EV profiles in inflammatory conditions. They support the idea that in inflamed joints with predominant neutrophil infiltration, neutrophil-derived EVs substantially contribute to the ensemble of SF-EVs. The lipidome analysis of EVs can form a first step in charting the cellular origin of the total SF EV population and defining a molecular lipid endotype of neutrophil-associated arthropathies, which can lead to the development of more precise disease diagnosis and treatment strategies.

## 5. Acknowledgments

We thank Jeroen Jansen for running the MS for the lipidomics experiment. We also thank Sander Tas from the Department of Rheumatology and Clinical Immunology, Amsterdam University Medical Centers, for supplying the synovial fluid material from their respective hospitals and Bernd Helms for critically reading the manuscript.

## 6. Declaration of Author Contributions

**Laura Varela:** Conceptualization, Formal analysis, Investigation, Visualization, Writing - Original Draft, Writing - Review & Editing; **Sanne Mol:** Conceptualization, Investigation, Resources, Writing - Review & Editing; **Esther W. Taanman - Kueter:** Investigation, Writing - Review & Editing; **Sarah E. Ryan:** Investigation, Resources; **Chris H.A. van de Lest:** Conceptualization, Software, Writing - Review & Editing; **Leonie S. Taams:** Resources, Writing - Review & Editing; **Esther C. de Jong:** Conceptualization, Resources, Writing - Review & Editing; **P. René van Weeren:** Conceptualization, Funding acquisition, Project administration, Supervision, Writing - Review & Editing; **Marca H.M. Wauben:** Conceptualization, Project administration, Supervision, Writing - Review & Editing.

## 7. Supplementary Material

**Supplementary Table 1:** Clinical and demographic parameters of patients with RA and SpA included.

| Patient | Diagnosis | Gender | Age (yrs) | Disease duration (yrs) | Rheumatoid factor | Medication |
| --- | --- | --- | --- | --- | --- | --- |
| 1 | SpA <sup>1</sup> | Male | 59 | 7 | - | NSAID*, anti-IL17 bDMARD** |
| 2 | SpA | Male | 52 | - | - | anti-TNF bDMARD |
| 3 | SpA | Female | 55 | 18 | - | NSAID, anti-TNF bDMARD |
| 4 | SpA | Female | 39 | 1 | - | DMARD***, B-complex vitamin |
| 5 | SpA | Male | 83 | 50 | Yes | DMARD, anti-TNF bDMARD |
| 6 | RA <sup>2</sup> | Female | 67 | 21 | Yes | NSAID |
| 7 | RA | Female | 85 | 10 | - | None |
| 8 | RA | Male | 36 | 18 | Yes | None |

<sup>1</sup> Spondyloarthritis
<sup>2</sup> Rheumatoid arthritis
\* Non-steroidal anti-inflammatory drugs
\*\* Biological disease-modifying antirheumatic drugs
\*\*\* Disease-modifying antirheumatic drugs

**Supplementary Figure 1:**
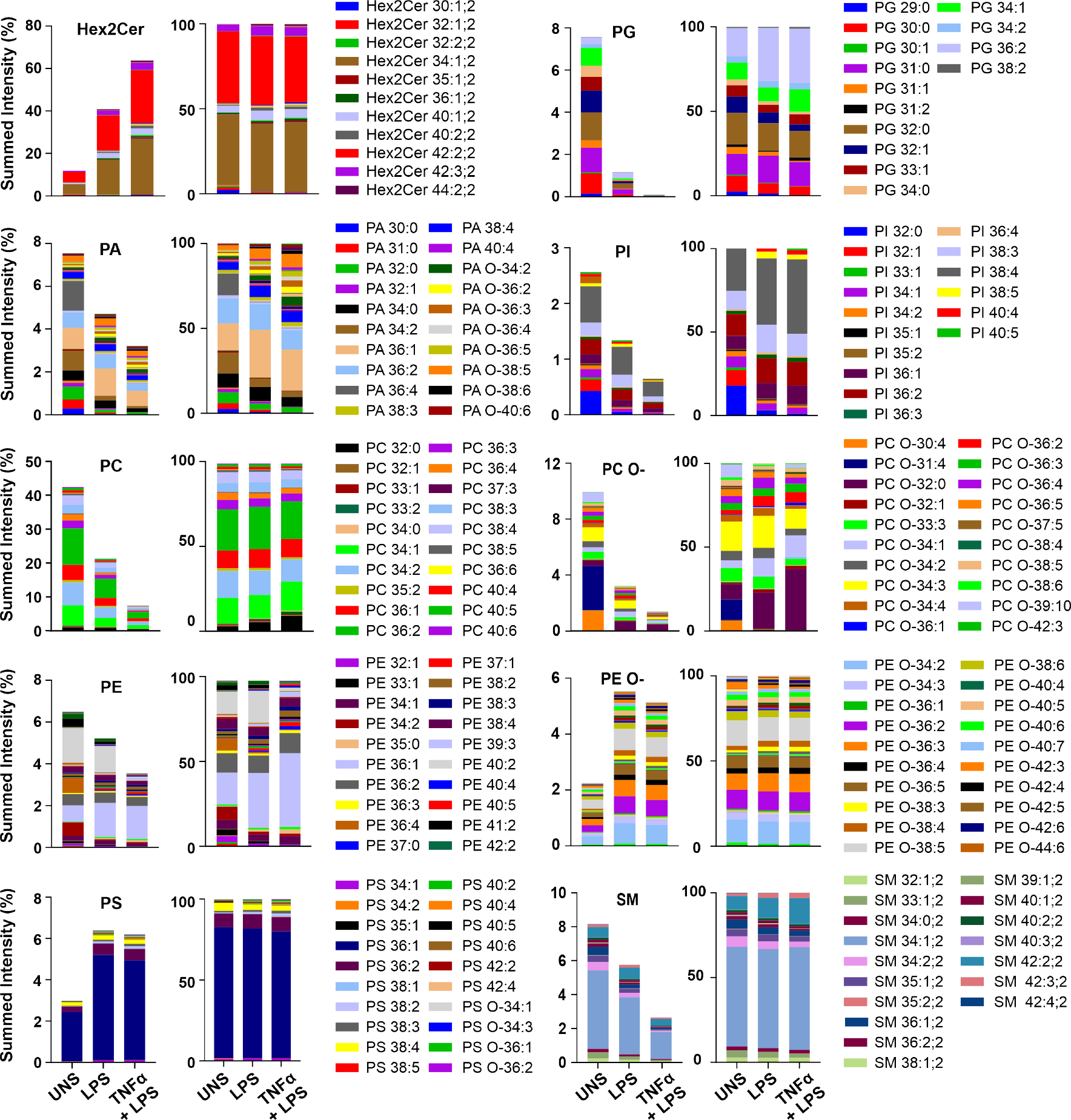
Composition of individual lipid species in Hex2Cer, PG, PA, PI, PC (O-), PE(O-), PS, and SM classes from Hu-nEVs. Lipid composition of the most abundant lipid classes from Hu-nEVs derived from unstimulated neutrophils (UNS; n = 5), single-stimulated neutrophils (LPS; n = 5), and dual-stimulated neutrophils (TNF*α*+LPS; n = 5). The left stacked bars graph of each class shows the number of lipid species in the overall lipidome. The bar graph immediately to the right displays the normalized amount of lipid species within the respective lipid class. Samples were normalized for each class and Hu-nEVs group by expressing the lipid intensity as a fraction of the sum of lipid intensities. Lipids were obtained from 100,000 xg purified Hu-nEVs with sucrose density gradients. Abbreviations: Hex2Cer (dihexosylceramide; lactosylceramide), PG (phosphatidylglycerol), PA (phosphatidic acid), O-(ether-linked lipids), PC (phosphatidylcholine), PE (phosphatidylethanolamine), PI (phosphatidylinositol), PS (phosphatidylserine), SM (sphingomyelin), Hu-nEVs (human neutrophil-derived extracellular vesicles).

**Supplementary Figure 2:**
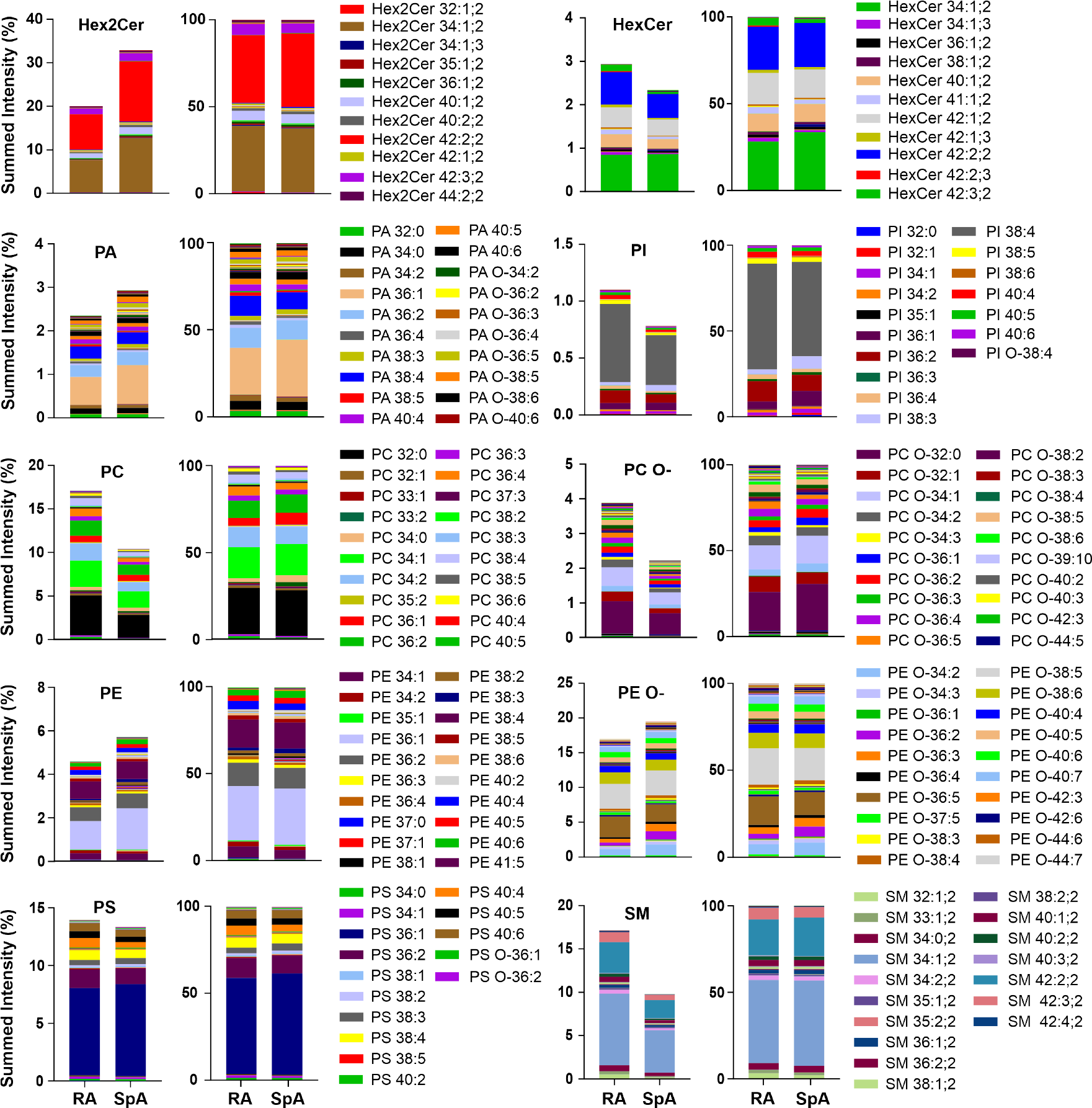
Composition of individual lipid species in Hex2Cer, HexCer, PA, PI, PC (O-), PE(O-), PS, and SM classes from SpA and RA SF-EVs. Lipid composition of the most abundant lipid classes of SF-EVs from RA (n = 3) and SpA (n = 5) patients. The left stacked bars graph of each class shows the amount of the lipid species in the overall lipidome. The bar graph to the right displays each class’s normalized amount of lipid species. Abbreviations: Hex2Cer (dihexosylceramide; lactosylceramide), HexCer (hexosylceramide; glucosylceramide), PA (phosphatidic acid), O-(ether-linked lipids), PC (phosphatidylcholine), PE (phosphatidylethanolamine), PI (phosphatidylinositol), PS (phosphatidylserine), SM (sphingomyelin), RA (rheumatoid arthritis), SpA (spondyloarthritis), SF-EV (Lipid composition of the most abundant lipid classes of SF-EVs from RA (n = 3) and SpA (n = 5) patients. The left stacked bars graph of each class shows the number of lipid species in the overall lipidome. The bar graph to the right displays each class’s normalized amount of lipid species. Abbreviations: Hex2Cer (dihexosylceramide; lactosylceramide), HexCer (hexosylceramide; glucosylceramide), PA (phosphatidic acid), O-(ether-linked lipids), PC (phosphatidylcholine), PE (phosphatidylethanolamine), PI (phosphatidylinositol), PS (phosphatidylserine), SM (sphingomyelin), RA (rheumatoid arthritis), SpA (spondyloarthritis), SF-EV (synovial fluid-derived extracellular vesicles).

**Supplementary Figure 3:**
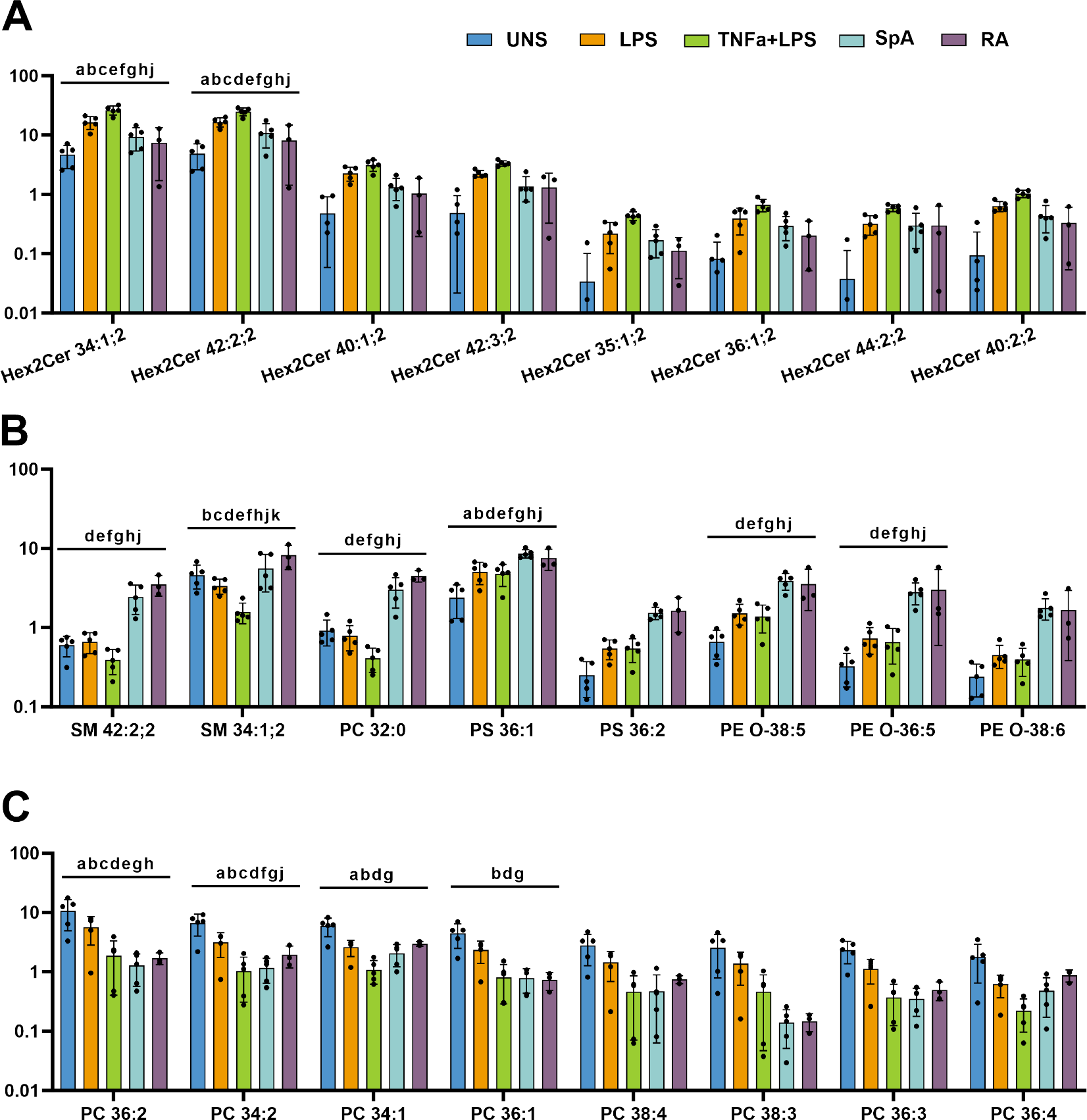
Bar plots of the concentration of individual lipid species that defined the loading plot of the PCA in Figure 3. Indicated as normalized lipid intensity as a percentage of the total lipid intensity. Shown are unstimulated neutrophil-derived EVs(UNS, n=5), single-stimulated neutrophil-derived EV samples(LPS, n=5), fully activated neutrophil-derived EVs (TNFa+LPS, n=5), SF-EVs derived from SpA patients(n=5), and SF-EVs derived from RA patients (n=3). Data are presented as bar plots with mean +/-standard deviation and the individual values as black dots. a = p < 0.05, UNS vs LPS; b = p< 0.05, UNS vs TNF*α* + LPS; c = p < 0.05, LPS vs TNF*α* + LPS; d = p < 0.05, UNS vs RA; e = p < 0.05, LPS vs RA; f = p < 0.05, TNF*α* + LPS vs RA; g = p < 0.05, UNS vs SpA; h = p < 0.05, LPS vs SpA; j = p < 0.05, TNF*α* + LPS vs SpA; k = p < 0.05, RA vs SpA. **A)** Concentrations of individual species where the SF-EV samples were above the unstimulated hu-nEV. **B)** Concentrations of individual species where the SF-EV samples were above all Hu-nEV. C) ConConcentrations of individual species where the SF-EVs samples were below the Hu-nEV or similar to the lowest, being the double-stimulated Hu-nEVs. Abbreviations: Hex2Cer (dihexosylceramide; lactosylceramide), O-(ether-linked lipids), PC (phosphatidylcholine), PE (phosphatidylethanolamine), PS (phosphatidylserine), SM (sphingomyelin), UNS (Unstimulated), TNF*α* (tumor necrosis factor-*α*) and lipopolysaccharide (LPS), Hu-nEVs (human neutrophil-derived extracellular vesicles), RA (rheumatoid arthritis), SpA (spondyloarthritis), SF-EV (synovial fluid-derived extracellular vesicles).

**Supplementary Figure 4:**
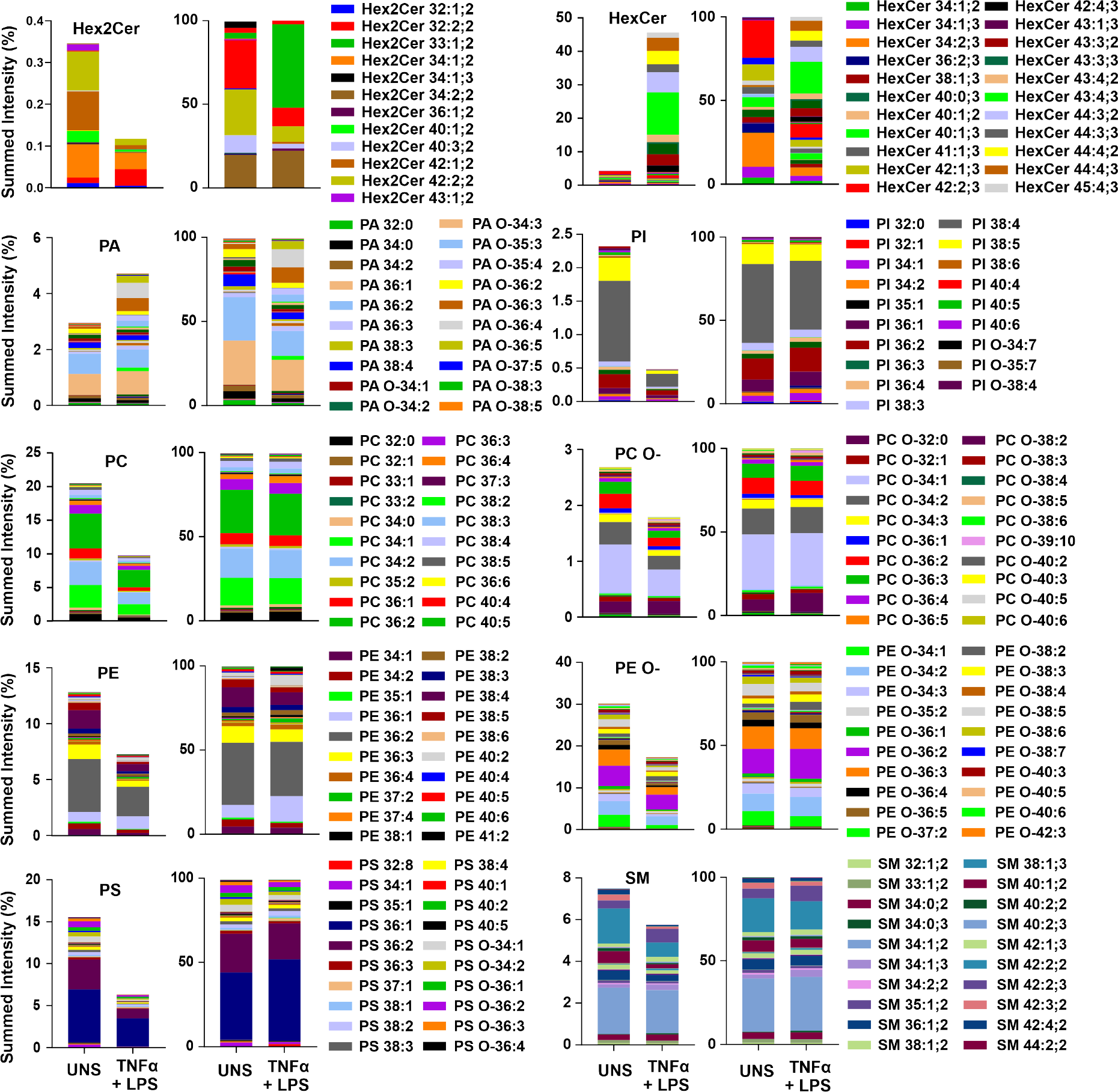
Composition of individual lipid species in Hex2Cer, HexCer, PA, PI, PC (O-), PE(O-), PS, and SM classes from Eq-nEVs. Lipid composition of the most abundant lipid classes from Eq-nEVs derived from unstimulated neutrophils (UNS; n = 3) and dual stimulated neutrophils (TNF*α* + LPS; n = 3) Eq-nEVs. The left stacked bars graph in each class shows the number of lipid species in the overall lipidome. The bar graph immediately to the right displays the normalized amount of lipid species in each class. Abbreviations: Hex2Cer (dihexosylceramide; lactosylceramide), HexCer (hexosylceramide; glucosylceramide), PA (phosphatidic acid), O-(ether-linked lipids), PC (ester-linked phosphatidylcholine), PE (phosphatidylethanolamine), PI (phosphatidylinositol), PS (phosphatidylserine), SM (sphingomyelin), Eq-nEVs (equine neutrophil-derived extracellular vesicles).

## References

Alarcon-Barrera, J.C., von Hegedus, J.H., Brouwers, H., Steenvoorden, E., Ioan-Facsinay, A., Mayboroda, O.A., Ondo-Mendez, A., Giera, M., 2020. Lipid metabolism of leukocytes in the unstimulated and activated states. Anal Bioanal Chem 412, 2353–2363. doi:10.1007/s00216-020-02460-8.

Benton, H.P., Want, E.J., Ebbels, T.M.D., 2010. Correction of mass calibration gaps in liquid chromatography–mass spectrometry metabolomics data. Bioinformatics 26, 2488–2489. URL: https://academic.oup.com/bioinformatics/article/26/19/2488/229139, doi:10.1093/bioinformatics/btq441.

Bert, S., Nadkarni, S., Perretti, M., 2023. Neutrophil-T cell crosstalk and the control of the host inflammatory response. Immunological Reviews 314, 36–49. URL: https://onlinelibrary.wiley.com/doi/abs/10.1111/imr.13162, doi:10.1111/imr.13162.

Bligh, E.G., Dyer, W.J., 1959. A rapid method of total lipid extraction and purification. Can. J. Biochem. Physiol. 37, 911–917. URL: http://www.nrcresearchpress.com/doi/10.1139/o59-099, doi:10.1139/o59-099.

Boere, J., van de Lest, C.H.A., Libregts, S.F.W.M., Arkesteijn, G.J.A., Geerts, W.J.C., Nolte-’t Hoen, E.N.M., Malda, J., van Weeren, P.R., Wauben, M.H.M., 2016. Synovial fluid pretreatment with hyaluronidase facilitates isolation of CD44+ extracellular vesicles. Journal of Extracellular Vesicles 5, 31751. URL: https://www.tandfonline.com/doi/full/10.3402/jev.v5.31751, doi:10.3402/jev.v5.31751.

Boff, D., Crijns, H., Teixeira, M.M., Amaral, F.A., Proost, P., 2018. Neutrophils: Beneficial and Harmful Cells in Septic Arthritis. International Journal of Molecular Sciences 19, 468. URL: https://www.mdpi.com/1422-0067/19/2/468, doi:10.3390/ijms19020468. number: 2 Publisher: Multidisciplinary Digital Publishing Institute.

Born, L.J., Khachemoune, A., 2023. Extracellular vesicles: a comprehensive review of their roles as biomarkers and potential therapeutics in psoriasis and psoriatic arthritis. Clinical and Experimental Dermatology 48, 310–318. URL: https://doi.org/10.1093/ced/llac108, doi:10.1093/ced/llac108.

Chatterjee, S., Balram, A., Li, W., 2021. Convergence: Lactosylceramide-Centric Signaling Pathways Induce Inflammation, Oxidative Stress, and Other Phenotypic Outcomes. International Journal of Molecular Sciences 22, 1816. URL: https://www.mdpi.com/1422-0067/22/4/1816, doi:10.3390/ijms22041816. number: 4 Publisher: Multidisciplinary Digital Publishing Institute.

Cokelaere, S.M., Plomp, S.G., de Boef, E., de Leeuw, M., Bool, S., van de Lest, C.H., van Weeren, P.R., Korthagen, N.M., 2018. Sustained intra-articular release of celecoxib in an equine repeated LPS synovitis model. European Journal of Pharmaceutics and Biopharmaceutics 128, 327–336. URL: https://linkinghub.elsevier.com/retrieve/pii/S0939641117309888, doi:10.1016/j.ejpb.2018.05.001.

Coletto, L.A., Rizzo, C., Guggino, G., Caporali, R., Alivernini, S., D’Agostino, M.A., 2023. The Role of Neutrophils in Spondyloarthritis: A Journey across the Spectrum of Disease Manifestations. Int J Mol Sci 24, 4108. URL: https://www.ncbi.nlm.nih.gov/pmc/articles/PMC9959122/, doi:10.3390/ijms24044108.

Couch, Y., Buzàs, E.I., Di Vizio, D., Gho, Y.S., Harrison, P., Hill, A.F., Lötvall, J., Raposo, G., Stahl, P.D., Théry, C., Witwer, K.W., Carter, D.R.F., 2021. A brief history of nearly EV-erything – The rise and rise of extracellular vesicles. Journal of Extracellular Vesicles 10, e12144. URL: https://onlinelibrary.wiley.com/doi/abs/10.1002/jev2.12144, doi:10.1002/jev2.12144.

Foers, A.D., Dagley, L.F., Chatfield, S., Webb, A.I., Cheng, L., Hill, A.F., Wicks, I.P., Pang, K.C., 2020. Proteomic analysis of extracellular vesicles reveals an immunogenic cargo in rheumatoid arthritis synovial fluid. Clin Transl Immunology 9, e1185. URL: https://www.ncbi.nlm.nih.gov/pmc/articles/PMC7648259/, doi:10.1002/cti2.1185.

Foers, A.D., Garnham, A.L., Chatfield, S., Smyth, G.K., Cheng, L., Hill, A.F., Wicks, I.P., Pang, K.C., 2021. Extracellular Vesicles in Synovial Fluid from Rheumatoid Arthritis Patients Contain miRNAs with Capacity to Modulate Inflammation. IJMS 22, 4910. URL: https://www.mdpi.com/1422-0067/22/9/4910, doi:10.3390/ijms22094910.

Gasser, O., Hess, C., Miot, S., Deon, C., Sanchez, J.C., Schifferli, J.u.A., 2003. Characterisation and properties of ectosomes released by human polymorphonuclear neutrophils. Experimental Cell Research 285, 243–257. URL: https://www.sciencedirect.com/science/article/pii/S0014482703000557, doi:10.1016/S0014-4827(03)00055-7.

Generali, E., Bose, T., Selmi, C., Voncken, J.W., Damoiseaux, J.G.M.C., 2018. Nature versus nurture in the spectrum of rheumatic diseases: Classification of spondyloarthritis as autoimmune or autoinflammatory. Autoimmunity Reviews 17, 935–941. URL: https://www.sciencedirect.com/science/article/pii/S1568997218301630, doi:10.1016/j.autrev.2018.04.002.

de Grauw, J.C., van de Lest, C.H., van Weeren, P.R., 2009. Inflammatory mediators and cartilage biomarkers in synovial fluid after a single inflammatory insult: a longitudinal experimental study. Arthritis Res Ther 11, R35. URL: http://arthritis-research.biomedcentral.com/articles/10.1186/ar2640, doi:10.1186/ar2640.

Groot Kormelink, T., Mol, S., De Jong, E.C., Wauben, M.H.M., 2018. The role of extracellular vesicles when innate meets adaptive. Semin Immunopathol 40, 439–452. URL: http://link.springer.com/10.1007/s00281-018-0681-1, doi:10.1007/s00281-018-0681-1.

Gu, Z., 2022. Complex heatmap visualization. iMeta 1, e43. URL: https://onlinelibrary.wiley.com/doi/abs/10.1002/imt2.43, doi:10.1002/imt2.43.

Gu, Z., Eils, R., Schlesner, M., 2016. Complex heatmaps reveal patterns and correlations in multidimensional genomic data. Bioinformatics 32, 2847–2849. URL: https://academic.oup.com/bioinformatics/article/32/18/2847/1743594, doi:10.1093/bioinformatics/btw313.

Hess, C., Sadallah, S., Hefti, A., Landmann, R., Schifferli, J.A., 1999. Ectosomes Released by Human Neutrophils Are Specialized Functional Units1. The Journal of Immunology 163, 4564–4573. URL:https://doi.org/10.4049/jimmunol.163.8.4564, doi:10.4049/jimmunol.163.8.4564.

Holopainen, M., Colas, R.A., Valkonen, S., Tigistu-Sahle, F., Hyvärinen, K., Mazzacuva, F., Lehenkari, P., Käkelä, R., Dalli, J., Kerkelä, E., Laitinen, S., 2019. Polyunsaturated fatty acids modify the extracellular vesicle membranes and increase the production of proresolving lipid mediators of human mesenchymal stromal cells. Biochimica et Biophysica Acta (BBA) - Molecular and Cell Biology of Lipids 1864, 1350–1362. URL: https://linkinghub.elsevier.com/retrieve/pii/S1388198119301131, doi:10.1016/j.bbalip.2019.06.010.

Iwabuchi, K., Masuda, H., Kaga, N., Nakayama, H., Matsumoto, R., Iwahara, C., Yoshizaki, F., Tamaki, Y., Kobayashi, T., Hayakawa, T., Ishii, K., Yanagida, M., Ogawa, H., Takamori, K., 2015. Properties and functions of lactosylceramide from mouse neutrophils. Glycobiology 25, 655–668. URL: https://academic.oup.com/glycob/article-lookup/doi/10.1093/glycob/cwv008, doi:10.1093/glycob/cwv008.

Iwabuchi, K., Nagaoka, I., 2002. Lactosylceramide-enriched glycosphingolipid signaling domain mediates superoxide generation from human neutrophils. Blood 100, 1454–1464. URL: https://doi.org/10.1182/blood.V100.4.1454.h81602001454_1454_1464, doi:10.1182/blood.V100.4.1454.h81602001454_1454_1464.

Jeucken, A., Molenaar, M.R., van de Lest, C.H., Jansen, J.W., Helms, J.B., Brouwers, J.F., 2019. A Comprehensive Functional Characterization of Escherichia coli Lipid Genes. Cell Reports 27, 1597–1606.e2. URL: https://linkinghub.elsevier.com/retrieve/pii/S2211124719304838, doi:10.1016/j.celrep.2019.04.018.

Kolonics, F., Kajdácsi, E., Farkas, V.J., Veres, D.S., Khamari, D., Kittel Merchant, M.L., McLeish, K.R., Lőrincz, M., Ligeti, E., 2021. Neutrophils produce proinflammatory or anti-inflammatory extracellular vesicles depending on the environmental conditions. Journal of Leukocyte Biology 109, 793–806. URL: https://doi.org/10.1002/JLB.3A0320-210R, doi:10.1002/JLB.3A0320-210R.

Lei, Q., Yang, J., Li, L., Zhao, N., Lu, C., Lu, A., He, X., 2023. Lipid metabolism and rheumatoid arthritis. Frontiers in Immunology 14. URL: https://www.frontiersin.org/articles/10.3389/fimmu.2023.1190607.

Leifer, V., Katz, J., Losina, E., 2022. The burden of OA-health services and economics. Osteoarthritis and Cartilage 30, 10–16. URL: https://linkinghub.elsevier.com/retrieve/pii/S106345842100738X, doi:10.1016/j.joca.2021.05.007.

Loh, J.T., Lam, K.P., 2022. Neutrophils in the pathogenesis of rheumatic diseases. Rheumatology and Immunology Research 3, 120–127. URL: https://www.degruyter.com/document/doi/10.2478/rir-2022-0020/html, doi:10.2478/rir-2022-0020. publisher: De Gruyter Open Access.

Lőrincz, M., Schütte, M., Timár, C.I., Veres, D.S., Kittel McLeish, K.R., Merchant, M.L., Ligeti, E., 2015. Functionally and morphologically distinct populations of extracellular vesicles produced by human neutrophilic granulocytes. Journal of Leukocyte Biology 98, 583–589. URL: https://doi.org/10.1189/jlb.3VMA1014-514R, doi:10.1189/jlb.3VMA1014-514R.

Lőrincz, M., Szeifert, V., Bartos, B., Szombath, D., Mócsai, A., Ligeti, E., 2019. Different Calcium and Src Family Kinase Signaling in Mac-1 Dependent Phagocytosis and Extracellular Vesicle Generation. Frontiers in Immunology 10. URL: https://www.ncbi.nlm.nih.gov/pmc/articles/PMC6928112/, doi:10.3389/fimmu.2019.02942. publisher: Frontiers Media SA.

Lőrincz, M., Timár, C.I., Marosvári, K.A., Veres, D.S., Otrokocsi, L., Kittel Ligeti, E., 2014. Effect of storage on physical and functional properties of extracellular vesicles derived from neutrophilic granulocytes. J Extracell Vesicles 3, 10.3402/jev.v3.25465. URL: https://www.ncbi.nlm.nih.gov/pmc/articles/PMC4275651/, doi:10.3402/jev.v3.25465.

Malda, J., Benders, K.E.M., Klein, T.J., de Grauw, J.C., Kik, M.J.L., Hutmacher, D.W., Saris, D.B.F., van Weeren, P.R., Dhert, W.J.A., 2012. Comparative study of depth-dependent characteristics of equine and human osteochondral tissue from the medial and lateral femoral condyles. Osteoarthritis Cartilage 20, 1147–1151. doi:10.1016/j.joca.2012.06.005.

McCoy, A.M., 2015. Animal Models of Osteoarthritis: Comparisons and Key Considerations. Vet Pathol 52, 803–818. doi:10.1177/0300985815588611.

Merola, J.F., Espinoza, L.R., Fleischmann, R., 2018. Distinguishing rheumatoid arthritis from psoriatic arthritis. RMD Open 4, e000656. URL: https://www.ncbi.nlm.nih.gov/pmc/articles/PMC6109814/, doi:10.1136/rmdopen-2018-000656.

Miao, H.b., Wang, F., Lin, S., Chen, Z., 2022. Update on the role of extracellular vesicles in rheumatoid arthritis. Expert Reviews in Molecular Medicine 24, e12. URL: https://www.cambridge.org/core/journals/expert-reviews-in-molecular-medicine/article/update-on-the-role-of-extracellular-vesicles-in-rheumatoid-arthritis/D63428836AB65F8EEC38A6302A0ABA32, doi:10.1017/erm.2021.33. publisher: Cambridge University Press.

Mol, S., Hafkamp, F.M.J., Varela, L., Simkhada, N., Taanman-Kueter, E.W., Tas, S.W., Wauben, M.H.M., Groot Kormelink, T., de Jong, E.C., 2021. Efficient Neutrophil Activation Requires Two Simultaneous Activating Stimuli. Int J Mol Sci 22, 10106. doi:10.3390/ijms221810106.

Mol, S., Taanman-Kueter, E.W.M., van der Steen, B.A., Groot Kormelink, T., van de Sande, M.G.H., Tas, S.W., Wauben, M.H.M., de Jong, E.C., 2023. Hyaluronic Acid in Synovial Fluid Prevents Neutrophil Activation in Spondyloarthritis. Int J Mol Sci 24, 3066. doi:10.3390/ijms24043066.

Moran, C.J., Ramesh, A., Brama, P.A.J., O’Byrne, J.M., O’Brien, F.J., Levingstone, T.J., 2016. The benefits and limitations of animal models for translational research in cartilage repair. Journal of Experimental Orthopaedics 3, 1. URL: https://doi.org/10.1186/s40634-015-0037-x,doi:10.1186/s40634-015-0037-x.

Nakayama, H., Nagafuku, M., Suzuki, A., Iwabuchi, K., Inokuchi, J.I., 2018. The regulatory roles of glycosphingolipid-enriched lipid rafts in immune systems. FEBS Letters 592, 3921–3942. URL: https://onlinelibrary.wiley.com/doi/abs/10.1002/1873-3468.13275, doi:10.1002/1873-3468.13275.

Oke, S.L., McIlwraith, C.W., 2010. Review of the economic impact of osteoarthritis and oral joint-health supplements in horses, pp. 12–16.

Pernes, G., Morgan, P.K., Huynh, K., Giles, C., Paul, S., Smith, A.A.T., Mellett, N.A., Veiga, C.B., Collins, T.J., Silva, T.M.D., Lee, M.K., Meikle, P.J., Lancaster, G.I., Murphy, A.J., 2023. An immune cell lipid atlas reveals the basis of susceptibility to ferroptosis. URL: https://www.biorxiv.org/content/10.1101/2023.02.10.528075v1, doi:10.1101/2023.02.10.528075. pages: 2023.02.10.528075 Section: New Results.

Phuyal, S., Skotland, T., Hessvik, N.P., Simolin, H., Øverbye, A., Brech, A., Parton, R.G., Ekroos, K., Sandvig, K., Llorente, A., 2015. The Ether Lipid Precursor Hexadecylglycerol Stimulates the Release and Changes the Composition of Exosomes Derived from PC-3 Cells *. Journal of Biological Chemistry 290, 4225–4237. URL: https://www.jbc.org/article/S0021-9258(19)47011-7/abstract, doi:10.1074/jbc.M114.593962. publisher: Elsevier.

Povoleri, G.A.M., Durham, L.E., Gray, E.H., Lalnunhlimi, S., Kannambath, S., Pitcher, M.J., Dhami, P., Leeuw, T., Ryan, S.E., Steel, K.J.A., Kirkham, B.W., Taams, L.S., 2023. Psoriatic and rheumatoid arthritis joints differ in the composition of CD8+ tissue-resident memory T cell subsets. Cell Rep 42, 112514. doi:10.1016/j.celrep.2023.112514.

Qu, F., Zhang, H., Zhang, M., Hu, P., 2018. Sphingolipidomic Profiling of Rat Serum by UPLC-Q-TOF-MS: Application to Rheumatoid Arthritis Study. Molecules 23, 1324. URL:http://www.mdpi.com/1420-3049/23/6/1324, doi:10.3390/molecules23061324.

Raggi, F., Bartolucci, M., Cangelosi, D., Rossi, C., Pelassa, S., Trincianti, C., Petretto, A., Filocamo, G., Civino, A., Eva, A., Ravelli, A., Consolaro, A., Bosco, M.C., 2023. Proteomic profiling of extracellular vesicles in synovial fluid and plasma from Oligoarticular Juvenile Idiopathic Arthritis patients reveals novel immunopathogenic biomarkers. Frontiers in Immunology14. URL: https://www.frontiersin.org/articles/10.3389/fimmu.2023.1134747.

Roberts, C.A., Dickinson, A.K., Taams, L.S., 2015. The Interplay Between Monocytes/Macrophages and CD4+ T Cell Subsets in Rheumatoid Arthritis. Frontiers in Immunology 6. URL: https://www.frontiersin.org/articles/10.3389/fimmu.2015.00571.

Scott, I.C., Whittle, R., Bailey, J., Twohig, H., Hider, S.L., Mallen, C.D., Muller, S., Jordan, K.P., 2022. Rheumatoid arthritis, psoriatic arthritis, and axial spondyloarthritis epidemiology in England from 2004 to 2020: An observational study using primary care electronic health record data. Lancet Reg Health Eur 23, 100519. URL: https://www.ncbi.nlm.nih.gov/pmc/articles/PMC9557034/, doi:10.1016/j.lanepe.2022.100519.

Skotland, T., Llorente, A., Sandvig, K., 2023. Lipids in Extracellular Vesicles: What Can Be Learned about Membrane Structure and Function? Cold Spring Harb Perspect Biol , a041415URL: http://cshperspectives.cshlp.org/content/early/2023/06/05/cshperspect.a041415, doi:10.1101/cshperspect.a041415. company: Cold Spring Harbor Laboratory Press Distributor: Cold Spring Harbor Laboratory Press Institution: Cold Spring Harbor Laboratory Press Label: Cold Spring Harbor Laboratory Press Publisher: Cold Spring Harbor Lab.

Skougaard, M., Ditlev, S.B., Stisen, Z.R., Coates, L.C., Ellegaard, K., Kristensen, L.E., 2022. Four emerging immune cellular blood phenotypes associated with disease duration and activity established in Psoriatic Arthritis. Arthritis Research & Therapy 24, 262. URL: https://doi.org/10.1186/s13075-022-02956-x, doi:10.1186/s13075-022-02956-x.

Skriner, K., Adolph, K., Jungblut, P.R., Burmester, G.R., 2006. Association of citrullinated proteins with synovial exosomes. Arthritis & Rheumatism 54, 3809–3814. URL: https://onlinelibrary.wiley.com/doi/abs/10.1002/art.22276, doi:10.1002/art.22276.

Smith, C.A., Want, E.J., O’Maille, G., Abagyan, R., Siuzdak, G., 2006. XCMS: Processing Mass Spectrometry Data for Metabolite Profiling Using Nonlinear Peak Alignment, Matching, and Identification. Anal. Chem. 78, 779–787. URL: https://pubs.acs.org/doi/10.1021/ac051437y, doi:10.1021/ac051437y.

Sokolove, J., Lepus, C.M., 2013. Role of inflammation in the pathogenesis of osteoarthritis: latest findings and interpretations. Therapeutic Advances in Musculoskeletal 5, 77–94. URL: http://journals.sagepub.com/doi/10.1177/1759720X12467868, doi:10.1177/1759720X12467868.

Symington, F.W., 1989. CDw17: a neutrophil glycolipid antigen regulated by activation. The Journal of Immunology 142, 2784–2790. URL: https://doi.org/10.4049/jimmunol.142.8.2784, doi:10.4049/jimmunol.142.8.2784.

Taitt, H.A., Balakrishnan, R., 2023. Spondyloarthritides. Immunology and Allergy Clinics of North America 43, 593–612. URL: https://www.sciencedirect.com/science/article/pii/S0889856122008980, doi:10.1016/j.iac.2022.10.001.

Tautenhahn, R., Böttcher, C., Neumann, S., 2008. Highly sensitive feature detection for high resolution LC/MS. BMC Bioinformatics 9, 504. URL: https://bmcbioinformatics.biomedcentral.com/articles/10.1186/1471-2105-9-504, doi:10.1186/1471-2105-9-504.

Team, R., 2020. RStudio: Integrated Development Environment for R. URL: http://www.rstudio.com/.

Varela, L., van de Lest, C.H., 2023. Lipidome profiling of human and equine neutrophil-derived extracellular vesicles and their potential contribution to the ensemble of synovial fluid-derived extracellular vesicles during joint inflammation. URL: https://public.yoda.uu.nl/dgk/UU01/2ABJW2.html, doi:10.24416/UU01-2ABJW2.

Varela, L., van de Lest, C.H.A., Boere, J., Libregts, S.F.W.M., Lozano-Andrés, E., van Weeren, P.R., Wauben, M.H.M., 2023. Acute joint inflammation induces a sharp increase in the number of synovial fluid EVs and modifies their phospholipid profile. Biochimica et Biophysica Acta (BBA) - Molecular and Cell Biology of Lipids 1868, 159367. URL: https://www.sciencedirect.com/science/article/pii/S1388198123000914, doi:10.1016/j.bbalip.2023.159367.

Vorobjeva, N.V., Chernyak, B.V., 2020. NETosis: Molecular Mechanisms, Role in Physiology and Pathology. Biochemistry Moscow 85, 1178–1190. URL: https://link.springer.com/10.1134/S0006297920100065, doi:10.1134/S0006297920100065.

Zhang, T., de Waard, A.A., Wuhrer, M., Spaapen, R.M., 2019. The Role of Glycosphingolipids in Immune Cell Functions. Frontiers in Immunology 10. URL: https://www.frontiersin.org/articles/10.3389/fimmu.2019.00090.

